# An evolutionary compass for detecting signals of polygenic selection and mutational bias

**DOI:** 10.1101/173815

**Authors:** Lawrence H. Uricchio, Hugo C. Kitano, Alexander Gusev, Noah A. Zaitlen

## Abstract

Selection and mutation shape genetic variation underlying human traits, but the specific evolutionary mechanisms driving complex trait variation are largely unknown. We developed a statistical method that uses polarized GWAS summary statistics from a single population to detect signals of mutational bias and selection. We found evidence for non-neutral signals on variation underlying several traits (BMI, schizophrenia, Crohn’s disease, educational attainment, and height). We then used simulations that incorporate simultaneous negative and positive selection to show that these signals are consistent with mutational bias and shifts in the fitness-phenotype relationship, but not stabilizing selection or mutational bias alone. We additionally replicate two of our top three signals (BMI and educational attainment) in an external cohort, and show that population stratification may have confounded GWAS summary statistics for height in the GIANT cohort. Our results provide a flexible and powerful framework for evolutionary analysis of complex phenotypes in humans and other species, and offer insights into the evolutionary mechanisms driving variation in human polygenic traits.

**Impact summary:** Many traits are variable within human populations and are likely to have a substantial and complex genetic component. This implies that mutations that have a functional impact on complex human traits have arisen throughout our species’ evolutionary history. However, it remains unclear how processes such as natural selection may have acted to shape trait variation at the genetic and phenotypic level. Better understanding of the mechanisms driving trait variation could provide insights into our evolutionary past and help clarify why it has been so difficult to map the preponderance of causal variation for common heritable diseases.

In this study, we developed and applied methods for detecting signatures of mutation bias (i.e., the propensity of a new variant to be either trait-increasing or trait-decreasing) and natural selection acting on trait variation. We applied our approach to several heritable traits, and found evidence for both natural selection and mutation bias, including selection for decreased BMI and decreased risk for Crohn’s disease and schizophrenia.

While our results are consistent with plausible evolutionary scenarios shaping a range of traits, it should be noted that the field of polygenic selection detection is still new, and current methods (including ours) rely on data from genome-wide association studies (GWAS). The data produced by these studies may be vulnerable to certain cryptic biases, especially population stratification, which could induce false selection signals. We therefore repeated our analyses for the top three hits in a cohort that should be less susceptible to this problem – we found that two of our top three signals replicated (BMI and educational attainment), while height did not. Our results highlight both the promise and pitfalls of polygenic selection detection approaches, and suggest a need for further work disentangling stratification from selection.

## Introduction

Natural selection and mutation shape variation within and between populations, but the evolutionary mechanisms shaping causal variation for human traits remain largely unknown. Studies of selection in humans have often focused on classic selective sweeps (Sabeti et al., 2002; Voight et al., 2006; Sabeti et al., 2006; Hernandez et al., 2011; Enard et al., 2014), but other processes such as stabilizing selection (Gilad et al., 2006; Sanjak et al., 2017b), polygenic adaptation (Turchin et al., 2012; Berg and Coop, 2014), background selection (Charlesworth, 1994; McVicker et al., 2009), negative selection (Boyko et al., 2008), and soft sweeps (Messer and Petrov, 2013; Schrider and Kern, 2017) may also play an important role in shaping human diversity. Methods to detect selection under these more complex models are needed if we are to fulfill the promise of genomics to explain the evolutionary mechanisms driving the distribution of heritable traits in human populations (Pritchard et al., 2010).

With the recent proliferation of paired genotype and phenotype data from large human cohorts, it is now feasible to test for polygenic selection on specific traits (Turchin et al., 2012; Berg and Coop, 2014; Yang et al., 2015; Robinson et al., 2015; Field et al., 2016). While these studies have argued that polygenic selection is likely to be an important determinant of variation in traits such as height and skin pigmentation (Berg and Coop, 2014), important questions remain about the evolutionary mechanisms that drive complex trait variation. In particular, most previous studies of selection on human complex traits have focused either on polygenic adaptation (Turchin et al., 2012; Berg and Coop, 2014; Field et al., 2016; Berg et al., 2017; Racimo et al., 2018) or stabilizing/negative selection (Yang et al., 2015; Zeng et al., 2018; Simons et al., 2018), and have not incorporated mutational bias (*i.e.*, the propensity for new mutations to be preferentially trait-increasing or preferentially trait-decreasing). To detect polygenic adaptation, studies have relied on genotype data from multiple populations to probe the frequency and linkage properties of trait-associated alleles as compared to a null based on genome-wide SNPs (Turchin et al., 2012; Berg and Coop, 2014; Racimo et al., 2018), or have used haplotype-based statistics and dense sequence data from a single population (Field et al., 2016). Negative selection has been investigated by comparing empirical data to null models of the relationship between frequency and squared effect sizes (Schoech et al., 2017) or linkage disequilibrium and per-SNP heritability (Gazal et al., 2017). These studies have argued that selection acts on many traits, and that both negative (Gazal et al., 2017; Schoech et al., 2017) and positive (Berg and Coop, 2014; Berg et al., 2017) selection act on complex traits such as height and BMI.

While each of these approaches has provided insights into the evolution of complex traits, a more comprehensive view of trait evolution will require methods that can account for pleiotropic selection and incorporate signals of both adaptive and deleterious selection processes simultaneously. Recent progress has been reported in accounting for higher dimensional trait spaces (Berg et al., 2017; Simons et al., 2018), but there is a need for models and inference tools integrating signatures of positive and negative selection on complex traits (Sanjak et al., 2017b). Indeed, a natural way to model polygenic adaptation is to view stabilizing selection as a null process, with punctuated changes in the fittest trait value (herein called the “optimal trait value” or “trait optimum”) driving brief periods of adaptation (Barton, 1986; Jain and Stephan, 2017). Mutational bias may also be an important contributor to the evolutionary dynamics of complex traits (Charlesworth, 2013), but has not been directly incorporated into recent empirical studies. If there is a bias in the direction of effect of *de novo* mutations, populations may carry less standing variation for alleles that alter the phenotype in one direction than the other, potentially altering the dynamics of future adaptation to changes in the trait optimum. Moreover, biases in mutation rate for selected traits may induce detectable patterns in the relationship between allele frequency and effect size, for example by driving an excess of trait-increasing mutations among young (low-frequency) alleles but not old alleles.

Here, we develop a powerful method for detecting differences between ancestral and derived allele effect sizes – which may be driven by selection or mutational bias – within a single population. We show that the polarization of GWAS summary statistics by their ancestral/derived state (which we refer to as an evolutionary compass) provides information about the evolutionary processes shaping trait variation. We propose a simple summary statistic of the relationship between effect sizes and allele frequency, and show that it is sensitive to both mutational bias and various models of selection. We apply our approach and find non-neutral signals that are consistent with selection and mutational bias in the genetic variation underlying BMI, educational attainment, Crohn’s disease, schizophrenia, and height. We develop a model-based inference procedure to disentangle mutational bias from selection, and show that both processes are necessary to explain the observed GWAS summary data. We then perform a replication study in the UK Biobank for our top three signals, and find that educational attainment and BMI replicate in this more homogeneous cohort, while height does not. We discuss implications of our findings for human evolutionary history and GWAS of biomedically relevant traits.

## Results

### An evolutionary compass for GWAS

Associations between genotypes and complex traits are usually reported with respect to a reference allele that is arbitrarily chosen, which can obscure the direction of effect of new mutations. When selection acts on traits, it favors the reproductive success or survival of individuals with particular trait values, implying that the fitness effect of new mutations may depend on both the sign and magnitude of their impact on a selected trait. Therefore, we choose ancestral alleles as the reference state and explore the relationship between derived allele effect sizes and derived allele frequency, which we encode in a *β*DAF plot (*i.e.*, a plot encoding the relationship between the mean value of effect sizes *β* estimated in a GWAS 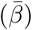 and derived allele frequency (DAF); Fig. 1). Note that we use estimated effect sizes from all alleles to compute 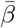, regardless of their significance.

**Figure 1:**
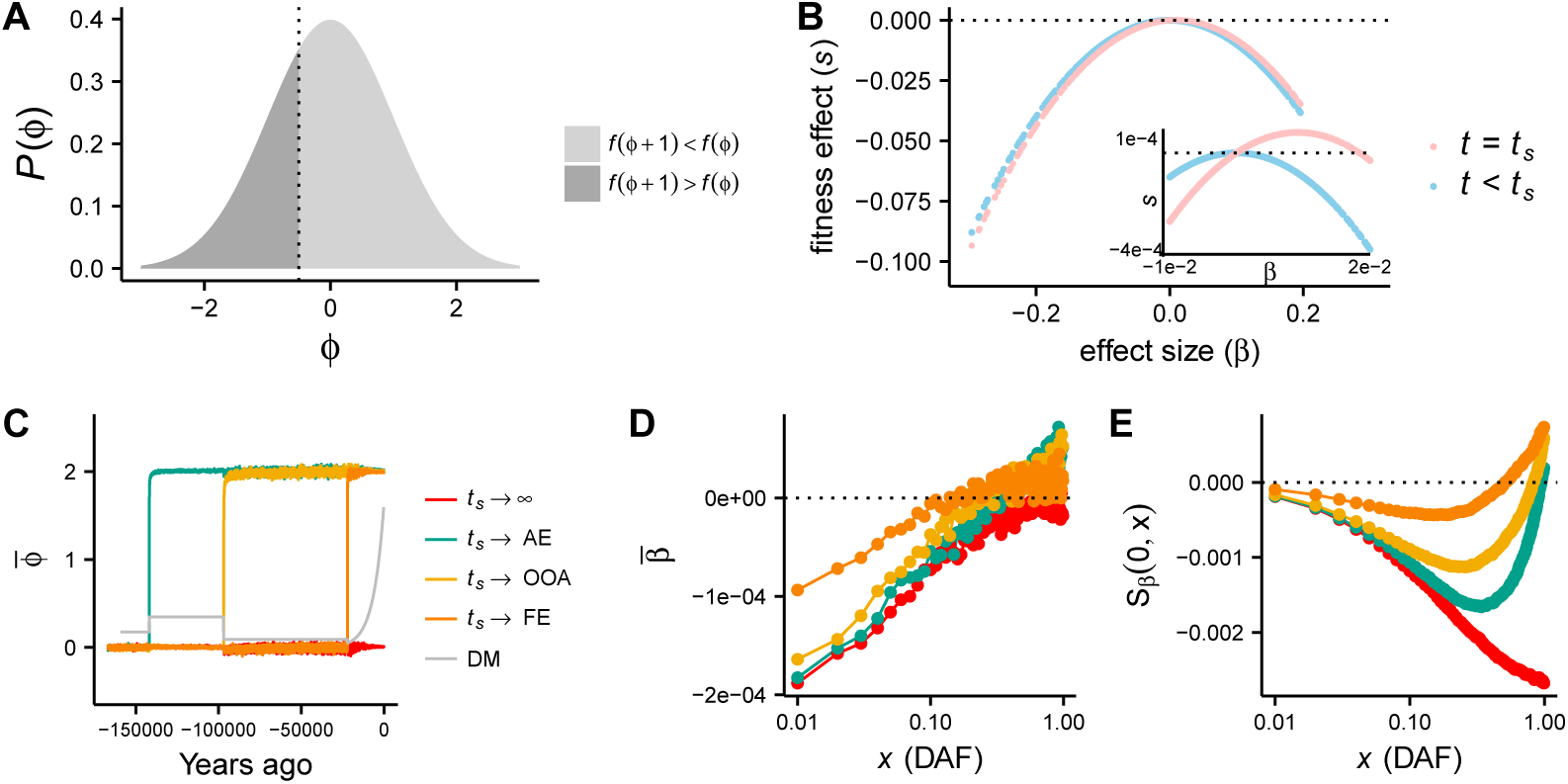
Panels A-B are schematics of the trait model, while C-E show simulation results. A: fitness impact of a *β* = 1 mutation, assuming a symmetric fitness function. At equilibrium, the trait distribution *P*(*ϕ*) is symmetric about the optimal value of the phenotype, *ϕ*_*o*_ = 0. The dashed line at 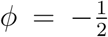 indicates the dividing line between individuals with increased fitness *f* after a *β* = 1 mutation (*f*(*ϕ* + 1) *> f*(*ϕ*))) from those with decreased fitness (*f*(*ϕ* + 1) *< f*(*ϕ*)). B: schematic of the relationship between effect size and fitness effect. At time *t* = *t*_*s*_, the optimal trait value *ϕ*_*o*_ increases, and trait-decreasing alleles have decreased fitness while trait-increasing alleles have increased fitness. Still, only trait-increasing alleles of small effect are on average fitness-increasing (inset). C. Mean trait value 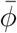 as a function of time for four simulated trait models, differentiated by the time of a shift in selection pressure. The simulated European demographic model is plotted in the background (not to scale) D. 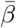 as a function of derived allele frequency (DAF) for each model simulated in C. Points represent the mean value of *β* computed over 100 independent simulations E. *S*_*β*_(0, *x*) as a function of DAF for each model plotted in C. D and E represent the mean over 100 independent simulations. (Abbreviations: *AE*: ancestral expansion, *OOA*: out-of-Africa, *FE*: founding of Europe, *DM*: demographic model).

A *β*DAF plot contains information about mutational bias and selection on the trait of interest. In the null case of a neutrally evolving trait for which trait-increasing and trait-decreasing mutations are equally likely, the *β*DAF curve will be flat will have expectation 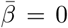 in all derived allele frequency bins. This is because in a neutral model, the probability with which an allele segregates at frequency *x* does not depend on effect size *β* (Fig. S2). A mutational bias towards trait-increasing or trait-decreasing alleles in the absence of selection on the trait will shift the mean value of *β* in the direction of the bias, but will not induce 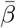 to depend on frequency (Fig. S2).

While many evolutionary processes will induce patterns in the *β*DAF plot (including directional selection, which we explore in the Supplemental Information with simulations and analytical calculations in Figs. S1-S3), we next consider an example of stabilizing selection with shifts in the optimal trait value for illustrative purposes. Stabilizing selection is typically parameterized by several parameters, but here we focus on the optimal phenotype value, *ϕ*_*o*_, which represents the value of the phenotype that confers the highest fitness. Applying a classic stabilizing selection model, we suppose that fitness is controlled by a Gaussian function centered at *ϕ*_*o*_ (Robertson, 1956; Barton, 1986; Simons et al., 2018) (Fig 1A). In addition, we suppose that trait-increasing mutations may be more or less likely than trait-decreasing mutations, which we capture with the parameter *δ* (defined as the proportion of trait-altering *de novo* mutations that increase trait values), and that *ϕ*_*o*_ can change, inducing a brief period of adaptation in which trait values within the population equilibrate to a new optimal value (Fig. 1B).

We performed forward simulations of complex traits under this model and explored how various evolutionary parameters affect the properties of a *β*DAF plot. Although we consider a very large range of possible parameter combinations when performing statistical inference in later sections, here we focused on 4 models, including one model of stabilizing selection in the absence of shifts in the optimal trait value, and three models that varied the timing of a shift (*t*_*s*_) in the optimal trait value *ϕ*_*o*_ (Fig. 1C). All of the models included a bias in mutation rate towards trait-decreasing alleles (*δ* = 0.4) and a European demographic model that was fit to patterns of European genomic diversity (Gravel et al., 2011). For models that include a shift in the optimal trait value, we considered a change (denoted ∆*ϕ*) equal to 2 standard deviations of the trait distribution. For reference, this would correspond to approximately a five-inch change in mean human height (Fryar et al., 2016). The remaining model parameters for the simulations in Fig. 1 are given in the Supplemental Information (page 26). To compute the mean effect size 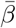 (Fig. 1D), we grouped alleles into 1% frequency bins and computed the mean effect size across all derived alleles within the bin.

When there is a bias towards trait-decreasing alleles and stabilizing selection acts on the trait, alleles at low frequencies have strongly negative effect sizes, which generates a positive correlation between effect size and frequency (Fig. 1D, red curves). When the optimal value of the phenotype increases in response to an environmental shift, the relationship between effect size and selection coefficient also transiently changes (Fig. 1B), driving some alleles with beneficial effects to higher frequencies (Fig. 1D, yellow, green, and orange curves). This effect transiently changes the relationship between allele frequency and effect size by promoting trait-increasing alleles to higher frequencies, generally increasing 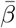 at all frequencies. Although 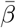 is always negative at low frequencies for the particular parameter combinations we investigated here, at high frequencies 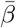 can become positive due to the preferential increase in frequency of trait-increasing alleles. These effects decay as the time since the shift event increases (Fig. 1D). The slow decay indicates that such patterns might be detectable in trait data for tens of thousands of years, with the time-scale for detection depending on the model parameters and the precision of effect size estimates in GWAS summary data.

### A statistical test for polygenic selection and mutational bias on oriented GWAS

Selection and mutational bias change the relationship between 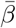 and DAF (Fig. 1; Figs. S1-S3). We desire a simple statistic that will differ from zero when selection and/or mutational bias act. We first consider the integral of the area under a *β*DAF plot, noting that while other statistics are also likely to be informative (Yang et al., 2015), not all choices will be robust to ancestral state uncertainty (see Supplemental Information). We approximate this integral with the sum (*S*_*β*_) over 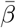 for derived allele frequency bins of some width *y*_*w*_.

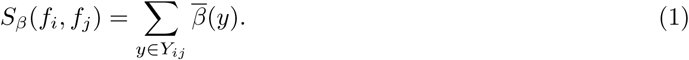

As in Fig. 1, we group alleles into frequency bins of width *y*_*w*_ such that the set *Y* is given by the sequence of frequency tuples *Y* = 〈(0, *y*_*w*_), (*y*_*w*_, 2*y*_*w*_),… (1–*y*_*w*_, 1)〉. *Y*_*ij*_ refers to the subsequence of *Y* that includes all elements indexed between *i* and *j*, and *f*_*i*_ is the lower frequency in the *i*-th tuple while *f*_*j*_ is the higher frequency in the *j*-th tuple. We choose *y*_*w*_ = 0.01 such that there are a large number of alleles within each bin.

The *S*_*β*_ statistic is sensitive to both mutational bias and selection, although these two distinct evolutionary processes drive distinct patterns in a *β*DAF plot. When mutational bias acts on new mutations in the absence of selection on the trait, there is no expectation of a relationship between the magnitude of effects and allele frequency, and hence the expected value of *β* is the same in all allele frequency bins (Fig. S2). This means that the expected value of *S*_*β*_ is simply equal to the mean effect size of new mutations multiplied by the number of bins, and that the sign of *S*_*β*_ is equal to the sign of the bias in mutational effects.

In the absence of mutational bias, *S*_*β*_ is sensitive to some (but not all) selection models. When a trait is under long-term stabilizing selection with no change in the optimal phenotype and no mutational bias, trait-increasing alleles are equally deleterious and equally likely to occur as trait-decreasing alleles, meaning that the expectation of *S*_*β*_ is 0 (which concords exactly with negative selection models – Fig. S3, purple curves). However, any bias in mutation rate towards trait-increasing or -decreasing alleles will drive *S*_*β*_ to have non-0 expectation (Fig. S3). In contrast to the neutral case with mutational bias, selection in conjunction with mutational bias causes the mean value of *β* to be greater in magnitude in bins of low derived allele frequency than high allele frequency, since selection will tend to constrain alleles with the largest effects to the lowest frequencies (Fig. 1C-E, red curves; Fig. S3).

The most interesting patterns emerge when stabilizing selection acts on a trait in conjunction with shifts in the fitness optimum. When stabilizing selection acts on a trait under along with mutational bias (*δ* = 0.4) and no shift in the optimum, *S*_*β*_(0, *x*) is negative at all *x*, and decreases as a function of frequency (red curve, Fig. 1E; we show in Fig. S3 and that this same pattern holds under a well-studied directional selection model for complex traits (Eyre-Walker, 2010)). When shifts in the optimum towards higher trait values occur, higher frequency variants have mean positive effect sizes, causing *S*_*β*_(0, *x*) to be non-monotonic and *S*_*β*_(0, 1) to potentially have positive sign. As time elapses since the shift, the high frequency trait-increasing alleles will tend to drift to the boundary and fix or be lost, causing this signal to gradually disappear (Fig. 1E). Note that mutational bias in the absence of selection can generate a non-zero *S*_*β*_ – however, mutational bias alone induces *S*_*β*_(0, *x*) to increase in magnitude monotonically and linearly in *x*, a pattern that is not expected for selection (Fig. S2 & Supplemental Information). We use model-based analyses to tease apart mutational bias and selection effects in later sections.

### A permutation procedure to generate a null distribution

While we have noted that *S*_*β*_ has an expected value of 0 under the neutral null without mutational bias, the variance of *S*_*β*_ depends on linkage between causal alleles and non-causals, since non-causal alleles will have non-0 estimated effect sizes when they are in LD with a causal allele. To control for potential confounding by LD, we developed a simple permutation-based procedure for computing the null-distribution of frequency-effect size relationships in GWAS summary statistics under a neutral evolutionary model. We first polarize all alleles such that the derived allele is the causal allele. Then, for each of 1,703 previously identified approximately independent linkage blocks (Berisa and Pickrell, 2016), we select a random sign (positive or negative with equal probability), and multiply all the effect sizes in the LD block by this sign. We then recompute the test statistic of interest, such as the correlation between frequency and MAF, on the randomized data. This generates a null distribution for the test statistic that conservatively accounts for the linkage between inferred effect sizes, and maintains the frequency spectrum and marginal distribution of the magnitude of inferred effect sizes.

### Application to GWAS summary data

We applied *S*_*β*_ within our framework to GWAS summary data for BMI, height (Wood et al., 2014), and educational attainment (Okbay et al., 2016) to assess its power for detecting selection, given that previous studies have suggested that these phenotypes may be under selection (Turchin et al., 2012; Berg and Coop, 2014; Robinson et al., 2015; Field et al., 2016; Racimo et al., 2018). We observe that effect sizes are correlated with frequency, and that *S*_*β*_ is a non-monotonic function of frequency (Fig. 2), consistent with selection and mutational bias (Fig. 1E; note that the large spike in both BMI and height at high frequency can be explained by ancestral uncertainty, Fig. S7). We reject the neutral null for all three traits (*p <*5e-4; Fig. 2 & Tab. 1). We then applied our method to six additional phenotypes, which we selected to span a wide range of phenotypes that we hypothesized might be targets of selection, including body size (Wood et al., 2014), psychiatric conditions (CDG Psychiatric Genomics Consortium, 2013), immune-related traits (Franke et al., 2010), reproductive traits (Day et al., 2015), and cardiovascular traits (Global Lipids Genetics Consortium et al., 2013). We find an additional non-neutral signal for Crohn’s disease, and a marginally significant signal for schizophrenia that narrowly missed a multiple testing correction (Tab. 1 & Figs. S8-16). When including only common variants in the test, we find that seven of nine phenotypes have *p*-values under 0.1, suggesting strong enrichment for selected traits among the test set despite failure for some of the tests to exceed a multiple testing correction (binomial *p*-value 3.0 *×* 10^−6^).

**Figure 2:**
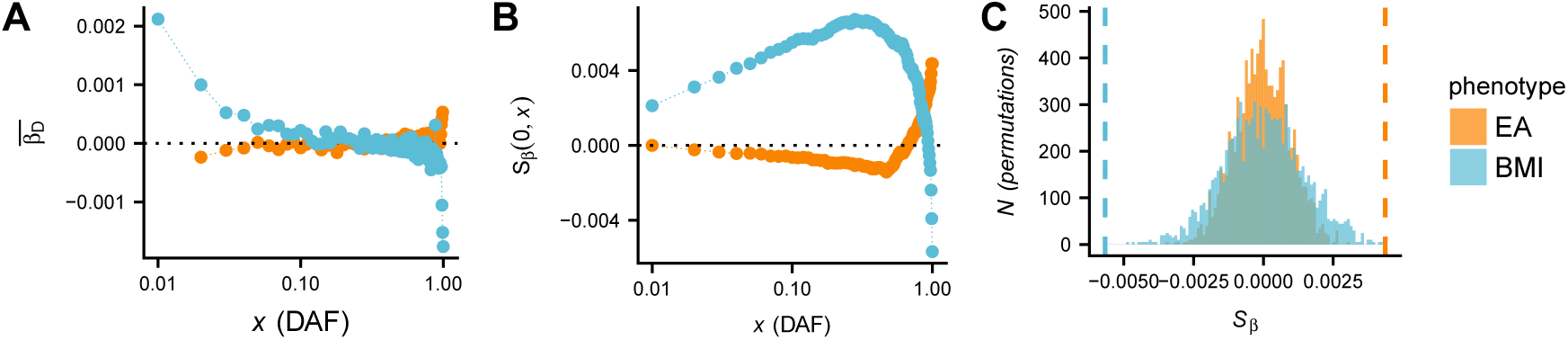
A: 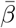 as a function of DAF for BMI and educational attainment (EA). B: *S*_*β*_(0, *x*) for the same data. C: neutral null distribution of *S*_*β*_(0, 1) obtained by permutations. The vertical dashed line indicates the observed value of *S*_*β*_(0, 1) in the GWAS summary data.

### Evolutionary models for selection on human traits

While our results show that *S*_*β*_ has a strong non-neutral signature for five of the nine phenotypes, we sought to further understand the evolutionary models that could explain these signals. Purifying selection alone seems an unlikely candidate for most of the phenotypes, because the sign of *S*_*β*_ is always the same as the sign of 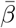 under a purifying selection model (Fig. S3 & Supplemental Information), a pattern violated by height, BMI, Crohn’s, and educational attainment. Models of directional selection for increased or decreased phenotype values in the absence of mutational bias also share this pattern, as do models of mutational bias in the absence of selection (Figs. S1-S2; Supplemental Information).

We hypothesized that these signals could be explained by a model of stabilizing selection, mutational bias, and shifts in the trait value conferring optimal fitness. We developed an evolutionary inference procedure based on rejection sampling (Tavaré et al., 1997) to infer the parameters that best fit the relationship between frequency and effect size that we observe in the data (see Supplemental Information). Briefly, we calculate *S*_*β*_(0.01, *x*) (scaled by *S*_*β*_(0.01, 0.99)) for a range of derived allele frequencies *x*, which we then use as summary statistics for rejection sampling. We remove the lowest frequency variants (*i.e.*, those with *x* < 0.01) for the purpose of this inference to avoid the potential impact of rare variant stratification on our results. We validated our method with extensive simulations, and found that it is a noisy estimator of the magnitude of these parameters (Fig. 3A&B – recall that *δ* corresponds to the proportion of new mutations that are trait-increasing), but has excellent power to estimate the direction of both mutational bias and shift in optimal trait value (Fig. 3C&D). Other parameters of the model (including heritability, polygenicity, effect size distribution, and the time of the shift in the fitness landscape; see Supplemental Information) were inferred with low accuracy as indicated by only modest correlations between inferred and true parameter values, indicating that the summary statistics we use contain little information about these parameters. Inferred distributions of these parameters therefore are not reported.

**Figure 3:**
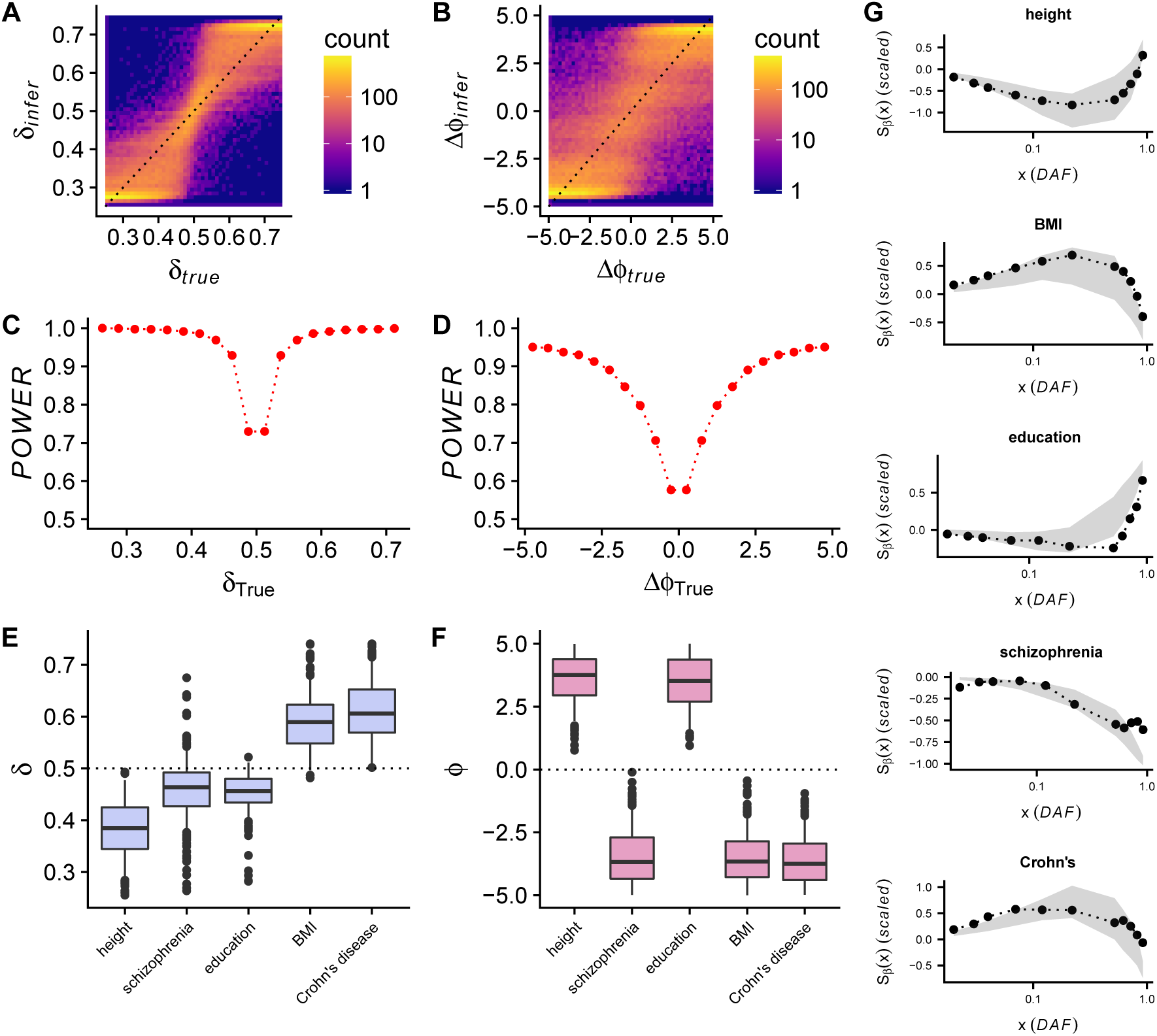
A-B: Inferred mutation bias (A) and selection shift (B) parameters as a function of true parameter values for our rejection sampling method. C-D: Power of our rejection sampling method to correctly identify the direction of mutation bias (C) and shift in optimal phenotype value (D), as a function of the true parameter value. E-F: Inferred approximate posterior distributions for five phenotypes that were identified as non-neutral. G: Out-of-sample simulations using parameters inferred in E-F, plotted with the data used to fit each model. Gray envelopes represent the middle 50% of simulation replicates, while the black points and curves show the observed data for each phenotype.

We find strong posterior support for a shift in optimal trait value (∆*ϕ*, measured in units of standard deviations of the population trait distribution) for all five phenotypes, as well as strong signals of mutational bias (*δ*). Data for height and educational attainment supported a shift towards increased trait values and a mutational bias towards mutations that decrease the phenotype, while Crohn’s disease and BMI supported shifts towards lower trait values and mutational bias towards trait-increasing alleles (*i.e.*, risk-increasing for Crohn’s). Schizophrenia data supported a shift towards lower risk, and a bias towards protective mutations, although a substantial minority of parameter estimates supported no mutational bias or a bias in the opposite direction (Fig. 3E). To further validate these findings, we resampled from the inferred parameter distributions and performed an independent set of forward simulations. We find that the summary statistics computed on these out-of-sample simulations (which were not used to fit the data) match the trends observed in our data, confirming that the modeling framework is capable of recapitulating the signals we observe in the data (Fig. 3G).

### Population stratification

Although *S*_*β*_ is not sensitive to confounding by ancestral uncertainty, we were concerned that these signals could potentially be explained by other confounders, such as uncorrected population stratification. While recent studies have suggested that methods to account for stratification in GWAS often over-correct (Field et al., 2016; Bulik-Sullivan et al., 2015), rare variant stratification remains especially difficult to account for in GWAS (Mathieson and McVean, 2012). We re-computed *S*_*β*_ on alleles with MAF*>* 1% and MAF*>* 5%, and found that the signals are robust within common alleles alone, suggesting that population structure is unlikely to confound our estimates (Figs. S8-16 and Tab. 1). Additionally, we recomputed *S*_*β*_ on GWAS from the UK Biobank (UKBB) for our top three signals, a replication cohort that employed strict population structure control in a more homogeneous group of samples. We replicate the signals for both educational attainment (*p* < 5*e −* 4) and BMI (*p* = 0.0095), noting that the educational attainment data in our two datasets contain some overlapping samples and hence are only partially independent and the BMI signal is somewhat weaker in UKBB than GIANT (Supplemental Information and Fig. S17). Interestingly, height was not replicated. We also applied two previous approaches that detected signals of selection acting on height when analyzing the GIANT summary statistics and found that neither replicated in the UKBB (Turchin et al. 2012; Yang et al. 2015; Fig. S6). These results are consistent with either over-correction of structure within the UKBB or under-correction in the GIANT data, although the latter seems more likely based on other recent studies (Berg et al. 2018; Sohail et al. 2018 – see Discussion).

## Discussion

Many studies have suggested that selection shapes human genetic variation (*e.g.*, Fay et al. 2001), and recent work has suggested that selection on complex traits may be a substantial driver of human adaptation (Hernandez et al., 2011). Here, we developed a novel empirical framework and a model-based rejection sampling approach for detecting polygenic selection and mutational bias that can be applied to GWAS summary data for a single population. We call this approach an “evolutionary compass”, because orienting alleles by their ancestral/derived status within our framework provides insight into the evolutionary processes shaping complex traits. We applied this evolutionary compass to GWAS summary data for nine phenotypes, and showed that five of them (educational attainment, height, Crohn’s disease, BMI, and schizophrenia) are consistent with a model of selection and mutation bias in shaping trait variation. Interestingly, among the top three signals that were uncovered with our method, height did not replicate in a more homogeneous cohort, while both BMI and educational attainment were replicated.

If selection acts on biomedically relevant complex traits such as Crohn’s disease and schizophrenia, there are important implications for the future of both medical and evolutionary genomics. In medical genomics, an ongoing debate about the genomic architecture of complex diseases is at the forefront of the field (Manolio et al., 2009). When strong selection acts on complex traits, it can elevate the role of rare alleles in driving trait variance (Lohmueller, 2014). If rare alleles contribute a larger fraction of the genetic variance than is expected under neutral models, then very large GWAS that use only array-based genotyping information are very unlikely to be able to capture these signals, and sequence-based studies and powerful rare variant approaches that are robust to evolutionary forces (including those not investigated here, such as partial recessivity) will be needed (Uricchio et al., 2016; Sanjak et al., 2017a; Hernandez et al., 2017). Moreover, recent work has suggested that the over-representation of Europeans in GWAS has limited the effectiveness of estimating polygenic risk scores in other human populations (Martin et al., 2017). This is problematic for the transfer of genomic research into the clinic, where precision medicine initiatives relying on personal genetic information will be most successful if genetic risk can be accurately predicted in diverse populations. While this inability to predict across populations could be driven by neutral demographic forces, if selection has driven numerous phenotypes to acclimate to local environmental conditions in ancestral human populations worldwide it could exacerbate this problem dramatically.

In the field of human evolutionary genomics, most studies have agreed that the impact of selection is widespread on the human genome, but the evolutionary mechanisms that drive genetic and phenotypic diversity have been widely debated (Hernandez et al., 2011; Enard et al., 2014; Schrider and Kern, 2017). In our study, we showed that GWAS summary statistics in Europeans for BMI and Crohn’s disease are consistent with a bias in mutation rate towards trait-increasing alleles, and a shift to a lower optimal value of the trait, while educational attainment is consistent with a mutational bias towards trait-increasing alleles and a shift towards higher values of the trait optimum. The signal for schizophrenia is consistent with an ancestral shift towards a lower optimum and a stabilizing selection, with or without a mutational bias. It should be noted that the model we used to fit these data assumed no more than one shift in the optimal phenotype value, whereas this quantity is likely to vary continuously with environmental conditions for some traits. Models that additionally account for sexual dimorphism (Stulp and Barrett, 2016), higher dimensional trait spaces (Simons et al., 2018), and evolutionary history of multiple populations (Berg and Coop, 2014; Racimo et al., 2018) may be required to better understand the generality of these results across human populations and traits.

Neanderthal introgression into modern humans has played an important role in shaping traits in non-African populations (Wall et al., 2013; McCoy et al., 2017). Given that Neanderthal alleles may contribute disproportionately to the genetic variance in some traits (Simonti et al., 2016) and that some high frequency trait-associated alleles have Neanderthal origins (Dannemann and Kelso, 2017; Prüfer et al., 2017), we hypothesized that Neanderthal alleles for traits under selection might show distinct patterns from modern human alleles. Although Neanderthal alleles do not share a common demographic history with modern human alleles, under the neutral null hypothesis we do not expect Neanderthal alleles to have an increasing or decreasing frequency/effect-size relationship, or to have a distribution that differs substantially from modern human alleles. We therefore computed *S*_*β*_ on alleles that have a Neanderthal origin (see Supplemental Information). Alleles for height and depression show strikingly different patterns than alleles with modern human origins (Fig. S18). The results for height can be explained by selection to promote Neanderthal height-increasing alleles to high frequency, either along the Neanderthal lineage predating human introgression, or after admixture with human populations. In contrast, our results for depression risk are consistent with an excess of depression risk from Neanderthals (Simonti et al., 2016), and selection preferentially driving large effect alleles to low frequency. We note that while Neanderthal alleles are not subject to the same biases in ancestral state uncertainty as modern human alleles, inferences of selection could still be biased by population stratification.

Inference of selection on complex traits is vulnerable to several possible confounders, including population stratification and pleiotropic selection on off-target phenotypes (Novembre and Barton, 2018). Although theory suggests that stratification should be straightforward to detect and correct at high frequency variants in large samples (Patterson et al., 2006), an uncorrected bias in the inferred *β* values due to population structure can make our test (as well most others, such as Berg and Coop 2014; Yang et al. 2015; Field et al. 2016; Berg et al. 2017; Racimo et al. 2018) anti-conservative. We used a series of experiments to show that population structure is unlikely to bias the majority of our results, including showing that the signal is robust to the exclusion of rare alleles and performing a replication study in an external cohort. However, one of our strongest signals (height) did not replicate the UK Biobank, while two other signals of selection suggested by earlier height studies also did not replicate in the UK Biobank (Turchin et al., 2012; Yang et al., 2015). The non-replication of height is in concordance with other recent studies finding reduced evidence for selection on height in the UK Biobank cohort applying different methods (Berg et al., 2018; Sohail et al., 2018). The most conservative interpretation of the non-replication of height is to suppose that the some of the signals we and others observed in the GIANT cohort are driven by population stratification, and the UK Biobank analysis correctly removes this spurious contamination. Further research is needed to better disentanlge stratification and selection, and caution in interpreting the results of polygenic selection tests is warranted while this field develops, even in more homogeneous cohorts such as the UK Biobank (Novembre and Barton, 2018).

Pleiotropy can also induce biases in complex trait selection detection. Selection on a trait that has correlated effect sizes with another trait could result in false positives, in which the neutral trait is spuriously identified. This is a general limitation of most complex trait selection methods (but see Berg et al. 2017; Simons et al. 2018). This phenomenon is clearly highlighted by our results on educational attainment, a phenotype which had no meaning until recent historical times. While it would be tempting to identify a cognition-related phenotype as the target of selection, it is possible that any trait with a cryptic shared genetic basis and correlated effect sizes could be the target, and that the timing of any such shift to larger trait values could have predated the human migration out-of-Africa. Thus, we cannot rule out a role for correlated phenotypes in driving these signals, and our results do not imply differences in phenotypes or polygenic scores between Europeans and any other group. More work on disentangling selection targets with a common genomic basis will be needed as the field progresses.

Among the nine traits that we tested, we found that four had strong non-neutral signals, and six of the nine had marginal evidence for non-neutrality (we have conservatively removed height, given its non-replication in the UK Biobank). However, this does not imply that the others are not subject to selection or mutation bias. The power of our test depends on the strength of selection, the polygenicity of the trait, the heritability of the trait, the mutational bias, and the amount of ancestral uncertainty, in addition to the size of the GWAS. If a trait is under strong stabilizing selection, but the mutation rate of trait-increasing and -decreasing alleles is exactly equal, then our test has no power. Since we rely on the polarization of alleles by ancestral state, increased uncertainty in ancestral state will also decrease our power (see Supplemental Methods). Moreover, if selection is weak, a small number of causal alleles drive variance in the trait, or the trait is only weakly heritable, power is greatly diminished. However, increased sample sizes in GWAS will increase power, because the variance of effect size estimates for even weak effect causal alleles decreases with sample size. In addition to working directly on GWAS summary statistics from a single population, one strength of our permutation-based approach is that other informative statistics, such as the absolute value of the deviation between ancestral and derived effect sizes, could also easily be applied, and may have higher power. In future studies, it will be advantageous to compare various summary statistics and to apply our approach to species undergoing rapid environmental changes.

**Table 1:** *P*-values corresponding to GWAS summary statistics for nine phenotypes that we hypothesized may be under selection. Values in the first column include all alleles, while the second and third columns correspond to tests including only alleles with MAF > 1% and MAF > 5%, respectively. The UK Biobank tests were performed on all alleles above 1% in frequency. **: Tests that pass a multiple testing correction, *p* < 0.005. *: Tests that were marginally significant (*p* < 0.05). *BMI: body mass index, WHR-BMI: waist-hip ratio adjusted for body mass index, GLL: global lipid levels*

| Phenotype | $S_\beta(0, 1)$ p-value | $S_\beta(0.01, 0.99)$ p-value | $S_\beta(0.05, 0.95)$ p-value | UKBB replication |
| --- | --- | --- | --- | --- |
| Height | < 0.0005** | < 0.0005** | < 0.0005** | 0.416 |
| BMI | < 0.0005** | < 0.0005** | < 0.0005** | 0.0095 |
| Education | < 0.0005** | < 0.0005** | < 0.0005** | < 0.0005 |
| WHR-BMI | 0.566 | 0.8325 | 0.341 | NA |
| GLL | 0.4655 | 0.434 | 0.5645 | NA |
| Crohn's disease | 0.0025** | 0.0075* | 0.018* | NA |
| Menopause onset | 0.1585 | 0.46 | 0.0475* | NA |
| Depression | 0.3915 | 0.01* | 0.0305* | NA |
| Schizophrenia | 0.0085* | 0.0035** | 0.0625 | NA |

## Materials and Methods

### Software

We wrote software in Python to implement the statistical test described herein, and we developed a custom simulator of demography and selection. The code to run our statistical test is freely available online ( https://github.com/uricchio/PASTEL). The simulations (and their validation) are described in the Supplemental Information.

## Data availability

We used publicly available GWAS summary data from each of the original studies of the phenotypes that we analyzed, including height, BMI, and BMI-WHR (Wood et al., 2014), schizophrenia and major depression (CDG Psychiatric Genomics Consortium, 2013), Crohn’s disease (Franke et al., 2010), menopause onset (Day et al., 2015), educational attainment (Okbay et al., 2016), and global lipid levels (Global Lipids Genetics Consortium et al., 2013). Post-processed data files (which map individual alleles to linkage blocks in the human genome) from each of these studies are available upon request, or could be generated by downloading the original GWAS summary data and mapping the alleles using the linkage data (Berisa and Pickrell, 2016). The UK Biobank summary data that we use in our replication study were obtained online and are also freely available (https://data.broadinstitute.org/alkesgroup/UKBB/).

## Acknowledgments

We acknowledge the support of NIGMS grant K12GM088033 and the Stanford IRACDA program (LHU). We thank our anonymous reviewers, Noah Rosenberg, Jeremy Berg, Doc Edge, Graham Coop, and Josh Akey for comments that substantially improved the manuscript. We also thank Jonathan Pritchard, Arbel Harpak, Yair Field, Ziyue Gao, and members of the Rosenberg, Coop, and Pritchard labs for helpful comments, as well as Josh Akey and Rajiv McCoy for providing a list of inferred Neanderthal alleles. We thank Nicole R. Gay for assistance in validating the software.

## Supplemental Information

### An evolutionary compass to detect selection and mutational bias

Selection on quantitative traits is known to induce dependency between mean squared effect sizes and derived allele frequency (DAF) (Eyre-Walker, 2010; Uricchio et al., 2016), but the relationship between effect sizes 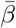 and DAF under selection models has not been widely explored. Under some models, for example when mutation rates for trait increasing and decreasing alleles are symmetric and the fitness cost of a mutation does not depend on the direction of its effect, selection will not induce a relationship between effect size and allele frequency. However, other models, such as those in which selection preferentially drives trait-increasing alleles to high frequency with selection pressure that increases as a function of *β*, may drive such a correlation (*e.g.*, Fig. S1A-C), which could potentially be detected in GWAS summary statistics data. We hypothesized that a better understanding of this relationship between *β* and DAF could provide new insights into the evolution of complex traits.

Models with symmetric mutation rates have often been applied to quantitative traits, but there is no *a priori* reason to suppose that this condition is met in natural populations. Indeed, it is possible that there are an unequal number of fixed bi-allelic sites genome-wide that can increase or decrease a phenotype relative to its current value, which naturally will change the relative rate of trait-increasing as compared to trait-decreasing alleles. For example, if past selection events have driven the phenotype to ever larger values, we might expect that the majority of trait-increasing alleles have already been fixed by positive selection in the evolutionary past, and that further recurrent mutations at these fixed sites would therefore decrease the phenotype.

Temporal variation in the optimal value of selected traits may also be an important determinant of the evolutionary dynamics of complex traits, as such changes may be a mechanism for polygenic adaptation (Jain and Stephan, 2017). When a population’s environment is altered, perhaps by migration, a change in climate, or the elimination/introduction of competing species, it is likely that selection pressures on phenotypes will also change. If the population persists long enough in this new environment it is expected that the phenotype mean will approach the new phenotype optimum, and the population will again be at equilibrium. In the intervening time, while the population is out of equilibrium, allele frequencies will shift as a function of their effect on the phenotype. Our goal is to capture selection’s impact on the causal variation during this out-of-equilibrium time period.

In this project, we sought to capture how mutational bias and selection pressure would affect the relationship between 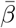 and allele frequency, and to use this information in designing a statistical test to detect mutational bias and selection. In the following sections, we use analytical theory and simulations to build intuition about the relationship between these evolutionary processes and patterns that could be observed in GWAS summary statistic data.

### Building intuition with the *β*DAF plot and *S*_*β*_

To build intuition about how evolutionary parameters affect the relationship between *β* and DAF we performed analytical calculations and simulations of simple evolutionary models that included directional selection, stabilizing selection, and mutational bias. Throughout these calculations, we suppose that although mutation rate can be biased towards trait-increasing or trait-decreasing alleles, the distribution of trait-increasing alleles is the same as that for trait-decreasing alleles.

#### Mutational bias alone

Mutational bias alone (*i.e.*, in the absence of selection) does not induce a dependency between 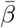 and DAF because mutations with differing *β*s are all equally likely to reach any given frequency. Hence, if the mutational bias is *δ* is the proportion of *de novo* mutations that are trait-increasing and the mean of the absolute value of a new mutation is 𝔼[|*β*|], then 𝔼[*β*] = *δ*𝔼[|*β*|] – (1 − *δ*)𝔼[|*β*|]. Note that when *δ* = 0.5, mutation rates are symmetric, and 𝔼[*β*] = 0.

Supposing that we decompose the frequency spectrum into *k* bins, when there is no selection 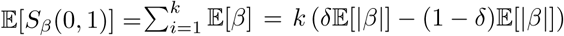. We perform this calculation for a variety of values of *δ* and a symmetric distribution of effect sizes, as well as stochastic simulations in which we select effect sizes randomly (Fig. S2), and observe excellent agreement between this simple calculation (solid lines) and the simulations (points).

𝔼[*S*_*β*_(0, *x*)] increases monotonically and linearly in *x* when mutational bias acts in the absence of selection on the trait, and the sign of *S*_*β*_ is always equal to the direction of the mutational bias (*i.e.*, mutational bias greater than 0.5 leads to positive *S*_*β*_, while mutational bias less than 0.5 leads to negative *S*_*β*_).

#### Polygenic adaptation

We next considered a toy model in which directional polygenic adaptation acts to increase the frequency of trait-increasing alleles. We suppose that trait-increasing alleles with effect size *β* have positive selection coefficients *s* with *β* = *s*^*τ*^, and that selection coefficients are drawn from a leptokurtic Γ-distribution peaked near 0. This model (which was proposed by Eyre-Walker for traits under negative selection (Eyre-Walker, 2010)) captures the idea that many alleles will have weak effects, and hence weak selective effects, while a small portion of alleles may have strong effects on the trait and be under relatively strong selection. We further suppose that the distribution and mutation rate of trait-increasing are the same as trait-decreasing alleles. We assume that the influence of binomial sampling is small and we do not account for it.

The time-dependent dynamics of the relationship between *β* and derived allele frequency are complex under such a model (because they depend on the transition probabilities from every possible initial frequency to every possible final frequency for all segregating selected alleles), but we can easily solve for the equilibrium state using diffusion theory. Under this model, the equilibrium frequency spectrum (i.e., the distribution of derived allele frequencies in the population at any given time) of trait increasing alleles is given by

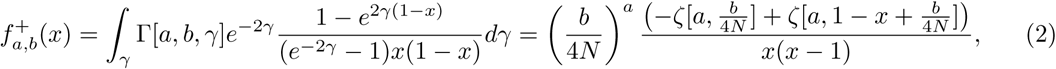

where 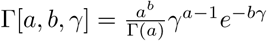, *a* and *b* are the parameters of the Γ-distribution, *ζ* is the Riemann zeta function, and *γ* is the population-scaled selection coefficient 2*Ns*. Trait-decreasing alleles have a neutral frequency spectrum, which is given by 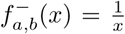. The mean value of *β* for trait increasing alleles at frequency *x* is given by

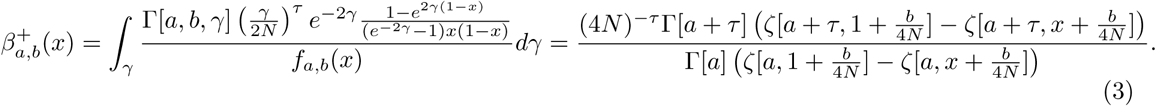

For trait-decreasing derived alleles, the mean value of *β* does not depend on the derived allele frequency *x* (because we assume trait-decreasing alleles evolve neutrally), and is given by

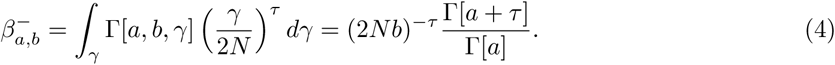

Combining terms and weighting by the frequency spectra for trait-increasing and trait-decreasing alleles, we find that the mean value of *β* as a function of DAF 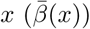 is given by

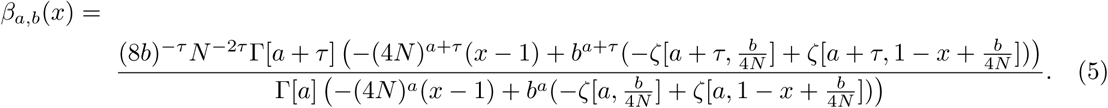

We performed stochastic simulations under this model, in addition to our calculations. For the simulations, we used a weak distribution of selective effects (𝔼[2*Ns*] ≈ 8) given by Γ[*a* = 0.0415, *b* = 0.00515625], which was inferred from human conserved non-coding polymorphism data (Torgerson et al., 2009). Our results show that trait-increasing alleles will be more likely to increase in frequency (Fig. S1B), and will outnumber trait-decreasing alleles at all frequencies, resulting in *β*_*a,b*_(*x*) > 0 at all frequencies (Fig. S1C). This implies that under this simple polygenic adaptation model, the *S*_*β*_ will increase monotonically in the direction of the selection (*i.e.*, if selection favors increases in trait values, *S*_*β*_(0, *x*) will be positive and monotonically increasing in *x*, while *S*_*β*_(0, *x*) will be negative and monotonically decreasing in *x* if lower trait values are preferred). Our analytical calculations from eqn. 5 (Fig. S1C, black curve) are in tight agreement with the simulations (Fig. S1C, points). Equations in this section were solved using *Mathematica*.

#### Mutational bias and negative selection

Negative selection against trait-altering alleles will also generate an increasing or decreasing *β*DAF relationship, if trait-increasing and trait-decreasing mutations are not equally likely to occur. To better understand this relationship quantitatively, we used analytical calculations under Eyre-Walker’s quantitative trait model to calculate the relationship between *β* and DAF.

We define mutational bias *δ* as the proportion of new mutations that are trait-increasing, and as in the previous section, we suppose that effect sizes are given by *β* = |*s*|^*τ*^. In contrast to the previous section (but in accordance with the original Eyre-Walker model), we suppose that all causal alleles are deleterious, regardless of their direction of effect. We suppose that *s* is drawn from a Γ-distribution.

Under this model, we are able to calculate *β*_*a,b*_(*x*) using the same procedure as in the previous section.
We find

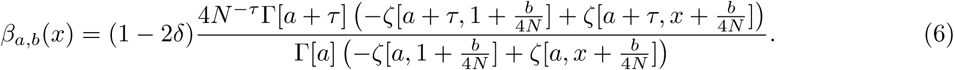

We performed stochastic simulations under this model for a variety of values of *δ*, again using a weak distribution of selection coefficients that was fit to human conserved non-coding sequences and *τ* = 0.5 (Torgerson et al., 2009). We find excellent agreement between our simulations and the analytical predictions (Fig. S3). When *δ* = 0.5, trait-increasing and trait-decreasing alleles are exactly balanced, and there is no dependency of *β* on DAF. In contrast, as *δ* becomes increasingly biased, trait-decreasing alleles further outnumber trait-increasing alleles (Fig. S3). Since larger effect alleles are subject to stronger selection, the difference is greater at low frequency than high frequency, generating a correlation between *β* and DAF. Taking *S*_*β*_(0, *x*) as the sum over the terms in eqn. 6, we find that *S*_*β*_ increases monotonically in magnitude in the same direction as the mutation bias under this model. In contrast to the mutational-bias only model, *S*_*β*_(0, *x*) does not increase linearly in magnitude as a function of *x* when selection acts.

#### Stabilizing selection with shifts

The previous sections show that a *β*DAF plot and *S*_*β*_ can capture signals of positive and negative selection as well mutational bias, but there is now substantial evidence that both positive and negative selection may act on complex traits. Indeed, it seems unlikely that polygenic adaptation will continue indefinitely over long evolutionary times, because this would imply that the trait would continue to increase or decrease over very long timescales. A more natural way to model negative selection punctuated with periods of adaptation is through stabilizing selection with shifts in the fittest value of the phenotype (Jain and Stephan, 2017).

We develop a polygenic selection quantitative trait model that maps selection coefficients *s* to effect sizes *β* using the well-established Gaussian stabilizing selection model (Robertson, 1956; Barton, 1986). We suppose that stabilizing selection acts on a trait, and that the fittest value of the trait is *ϕ*_*o*_ (also referred to as the “trait optimum”), such that the fitness *f* of an individual with trait value *ϕ* is given by

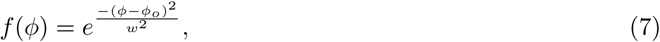

where *w* is the breadth of the fitness function. We additionally suppose that the trait *ϕ* has a normal distribution such that

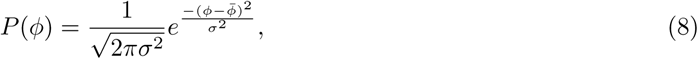

where *σ* is the standard deviation of the fitness distribution and 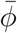 is the mean trait value in the population. Under these conditions, it is possible to solve for the per-generation, per-individual selection coefficient *s* as a function of the above model parameters for causal alleles of effect size *β*. We calculate *s* by marginalizing the fitness effect of a new mutation of effect size *β* across all fitness backgrounds. Our calculation proceeds similarly to previous work (Barton, 1986; Simons et al., 2018), although we retain the dependence on the current mean phenotype value 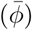 and the phenotype optimum (*ϕ*_*o*_), rather than assuming that the phenotype distribution is centered at the optimum. While the full expression for the expected change in allele frequency is provided in the next section, we note that when the trait is at equilibrium such that 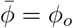,

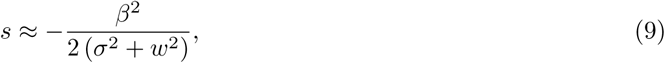

and the expected frequency change *δp* for an allele at frequency *p* with effect size *β* when 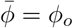 is

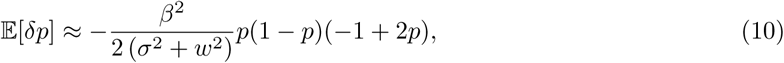

which implies that the trait evolves as if underdominant when the mean population trait value is centered close to the optimum.

#### Recasting the selection coefficient in terms of the trait distribution

To calculate the time-dependent selection coefficient *s*(*t*) of a site with effect size *β*, we first develop some results that allow us to recast the selection coefficient *s* at equilibrium as a function of the proportion of the population in which an allele of effect size *β* is fitness-increasing. Since the trait is normally distributed, the probability that an individual has phenotype value *ϕ*, *P* (*ϕ*), is given by 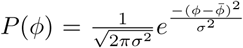. To calculate the selection coefficient for an allele with effect size *β*, we then marginalize across all trait backgrounds and possible genotypes in the population to obtain the mean fitness effect, since the fitness effect varies across trait backgrounds according to *f*(*ϕ*). The three possible genotypes for a causal variable allele are denoted as *aa* (homozygous ancestral), *ad* (heterozygous), and *dd* (homozygous derived). For each genotype, we calculate the fitness *w* of an allele at frequency *p* as

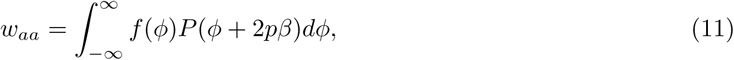

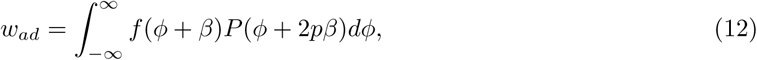

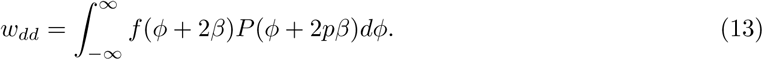

substituting in *P*(*ϕ*) and *f*(*ϕ*), these integrals can be solved to obtain 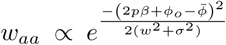, 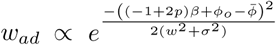, and 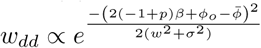. We then can solve for the expected change in frequency 𝔼[*δp*] at time of an allele at frequency *p* as

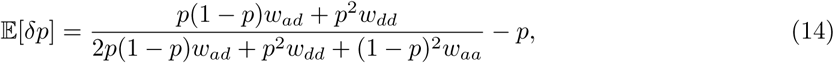

Taking a series expansion about *β* = 0, we find that

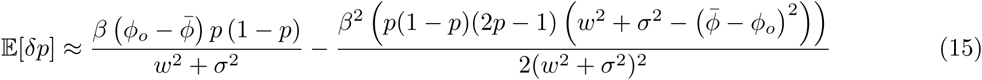

When 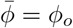, alleles evolve as if underdominant because 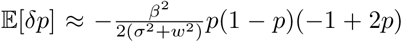. When 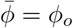 alleles may either be expected to increase in fitness (*i.e.*, be transiently positively selected) or decrease in frequency, depending on their effect size and the current mean population trait value 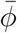. In our Wright-Fisher simulations, we replace the expected frequency change with eqn. 15 and track the current mean population trait value as a time-dependent quantity (see next section for more details).

#### Validation of simulations

We developed a custom simulator of our model in Python. Our simulator accommodates changes in population size, including explosive growth and bottlenecks, as well as any arbitrary distribution of selection coefficients. We perform a burn-in period of 5*N* generations for a simulated population of size *N*, during which we suppose that the population is close to equilibrium and hence *s* does not vary each generation. After the first burn-in, we perform an additional 5*N* generations of burn-in during which we recalculate 𝔼[*δp*] as a function of *β*, 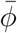, and *ϕ*_*o*_ in each generation with eqn. 15. To calculate the current value of 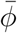, we sum over all *j* causal alleles, such that 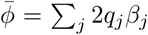, where *q*_*j*_ is the frequency of a site with effect size *β*_*j*_.

For simulations of polygenic adaptation, at some time *t*_*s*_ during the simulations, we reset *ϕ*_*o*_ to a new value, which induces the trait distribution to be out-of-equilibrium with respect to the fitness function. During the out-of-equilibrium period, a portion of the causal sites will be positively selected (specifically those that are fitness increasing when marginalizing across all trait backgrounds, as described in the previous section), while the remaining sites will be fitness-decreasing. Each generation, we recalculate *δp* based on the current configuration of 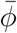 and *ϕ*_*o*_.

To validate our simulator, we simulated a complex selection and demographic model using previously published models of European demographic history (Gravel et al., 2011) and selection on human conserved elements (Torgerson et al., 2009) in SFS CODE (Hernandez, 2008) and compared the SFS CODE frequency spectrum to the frequency spectrum in our simulations. When no shift in the trait optimum occurs, selection coefficients in our model have the same expected value as in the standard Wright-Fisher model with underdominance, so the frequency spectra should be similar. However, since underdominance is not straightforward to simulate in SFS CODE, for this set of validation simulations we replaced the underdominance term in our simulations with the standard genic selection model. Results of these simulations are plotted in Fig. S4. We observe good agreement between the two spectra overall, although our model results in a slight over-representation of rare alleles and a slight under-representation of common alleles relative to SFS CODE. Note that there is weak LD in the SFS CODE simulations and that our selection model differs slightly, so we do not expect perfect agreement. For the SFS CODE simulations, we used the following command line: sfs code 1 500 -N 1000 -n 100 -A -t 0.001 -r 0.0 -TE 0.405479 -Td 0 P 0 1.982738 -Td 0.265753 P 0 0.128575 -Td 0.342466 P 0 0.554541 -Tg 0.342466 P 0 55.48 -L 100 150 -l g 0.5 R -a N R -W 2 0 1 1 0.0415 0.00515625 -s <random seed> -o <out>

For the simulations presented in Fig. 1, the curves represent the mean over 100 independent simulations. We used *N* = 7, 000 (where *N* is the ancestral population size) for the simulations in Fig. 1., but rescaled the population size for computational efficiency to 2,000 for parameter inference (see next section). The simulation code was implemented in Python and numpy, and will be freely available and posted on Github.

#### *S*_*β*_ in simulation

We performed simulations of Gaussian stabilizing selection with shifts in the optimal phenotype value for a wide range of parameter combinations. Since we ultimately apply our results to data from European GWAS, we apply a European demographic model (including ancestral expansion and bottleneck events, as well as recent exponential growth) that was fit to patterns of genomic diversity (Gravel et al., 2011). Our goal in these simulations is to understand the qualitative behavior of *S*_*β*_ as a function of evolutionary parameters (Fig. 1C-E), and in subsequent sections we develop a more quantitative approach for mapping evolutionary parameters to observed patterns in GWAS summary statistics.

For simulations shown in Fig. 1, we used *N* = 7, 000 (in accordance with (Gravel et al., 2011)), a mutational bias of *δ* = 0.4, *h*^2^ = 0.8, and selection strength *ω* = 1. Effect sizes drawn from a Gamma distribution with *a* = 0.037 and *b* = 0.0008, and assumed a genome-wide polygenicity of 0.02 (*i.e.*, 2% of target sites have a non-zero effect on phenotype). While these choices are arbitrary, we use rejection sampling to find parameter combinations that fit the observed data in the subsequent section.

On the advice of a reviewer, we also explored more gradual and less dramatic shifts than those we simulated in Fig. 1. We simulated shifts of 0.5, 1, and 2 trait standard deviations that occur linearly over 100 generations (*e.g.*, the optimum increases by 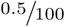 per generation in the 0.5 case), but otherwise with the same parameters as above. We performed this set of experiments to confirm that the dynamics we observe are not driven primarily by unrealistically abrupt shifts that would not be observed in nature. We found that a gradual shift with Δ*ϕ* = 2 had nearly identical patterns to the instantaneous case presented in Fig. 1 (Fig. S19A-C). However, reducing the total magnitude to 1 and 0.5 trait standard deviations drives substantial decreases in the magnitude of the signal (Fig. S19D-I). As expected, these results imply that smaller and more ancient shifts will be more difficult to detect, but instantaneous shifts drive similar patterns to gradual shifts (at least in this part of the parameter space).

#### Inferring evolutionary parameters

We developed a rejection-sampling based approach to infer evolutionary parameters (Tavaré et al., 1997). Rejection sampling uses informative summary statistics in model-based simulations to infer an approximate posterior distribution of model parameters. Simulations are performed under the model with parameters drawn from a wide range of possible combinations, and then the subset of simulations that most closely match the observed data are retained to build the posterior. The remaining simulations are rejected.

We performed simulations of Gaussian stabilizing selection with shifts in the optimal phenotype value using parameter values that were sampled from broad prior distributions, and we fit these simulated data to the observed *S*_*β*_(0.01, *x*) data for our phenotypes (we exclude alleles below 1% frequency to avoid the potential effects of rare variant stratification). As summary statistics, we take *S*_*β*_(0.01, *x*) for *x* ∈ {0.02, 0.03, 0.04, 0.07, 0.12, 0.22, 0.52, 0.62, 0.72, 0.82, 0.92}. Additionally, since the absolute magnitude of *S*_*β*_ depends on the units of the effect sizes, we defined scaled *S*_*β*_(*x*_*i*_, *x*_*f*_) as 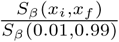, and use the scaled summary statistics as input to our method. As a final summary statistic, we take 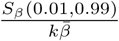, where 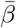 is the mean effect size over all effect sizes in the study (or simulation) and *k* is the number of frequency bins (*k* = 100 here). As we showed earlier in the Supplemental Information, *S*_*β*_ has expectation 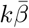 when only mutational bias acts on the trait, so this summary statistic has expectation 1 when only mutational bias acts on the trait.

We performed 10^5^ simulations, drawing evolutionary parameters from very wide prior distributions. Parameters of the model included heritability (*h*^2^, uniform from 0.5 to 1), shift in optimum phenotype (∆*ϕ*, uniform on -5 to 5 in units of the standard deviation of the ancestral trait distribution), polygenicity (Ψ, chosen uniformly from 0.004 to 0.02 of sites genome-wide being causal), effect size distribution parameters (chosen from the square root of a Γ-distributed according to *a* and *b*, with *a* chosen to be within a factor of 4 greater or less than 0.04 and *b* chosen to be within a factor of 4 of 8e-5 – this enforces selection coefficients to be drawn from a Γ-distribution), mutational bias *δ* (uniform on 0.25 to 0.75), the time of the shift in optimal phenotype (*t*_*s*_, uniform from -1 to 0.4 in coalescent units, corresponding to the time period ranging from 500,000 to 17,500 years ago), and ancestral uncertainty (unifom from 0.7 to 1, where 1 represents perfect ancestral assignment). Because we vary both heritability and the effect size distribution, we did not additionally vary the selection parameter of Gaussian stabilizing selection *ω*, which we set at 1. We rescaled the population size to *N* = 2 × 10^3^ for computational efficiency (the ancestral population size in the demographic model is *N* = 7 × 10^3^). Although we do not directly vary the selection coefficient distribution (because selection coefficients depend on effect sizes and the current trait variance *σ*^2^, and we sample effect size parameters as opposed to selection parameters), our simulations include both very weak selection per site and strong selection per site. After each simulation finishes, we mix the causal alleles with neutral alleles that are simulated under the same demographic model, and have effect size 0, representing the portion of the genome that is not causal for the trait but is included in the GWAS. We calculate the sum of squared differences between our simulated and observed statistics for all 10^5^ simulations, and retain the 0.25% of simulations that minimize this statistic.

To test our rejection sampling method, we masked the input parameter set of each simulation and attempted to infer the parameter values that were used to run the simulation. We used the maximum *a posteriori* estimate as a point estimate of the parameter value for these experiments. We found high predictability of the direction of both mutational bias *δ* and ∆*ϕ* (Fig. 3C&D), and noisy estimation of the magnitude of these parameters. Most other parameters were poorly predicted. We found that the true timing of selection events was highly correlated with the inferred timing, but a large number of outlier estimates make the estimates of this parameter less trustworthy, so we do not report it. All other parameters were inferred with low accuracy, hence we only report *δ* and ∆*ϕ*.

In addition to the power to detect mutation rate bias (*δ*) and shifts in optimal phenotype value (∆*ϕ*), we also considered the false positive rate for detection of these parameters (Fig. S5). We examined the subset of simulations among our 10^5^ that had |∆*ϕ*| < 0.25 and the subset with 0.45 *< δ* < 0.55 and assessed the proportion of these simulations for which we estimated maximum *a posteriori* parameters larger than *y* for all estimated values *y*. We found that the false positive rate is high for both small mutation rate biases and weak shifts, but low for effects as large as those we infer in the data, implying that our method is unlikely to result in a multiple false positives for shifts and mutation rate biases. Note that this null is very conservative, because all of the simulations we used to assess false positive rate include stabilizing selection, and the *δ* false positive simulations include shifts in selection while the ∆*ϕ* false positive simulations include bias in mutation rate.

It should be noted that while our empirical selection detection method incorporates linkage, our rejection sampling method does not. Linkage between causal alleles and non-causals could cause the data to have larger deviations from 0 in *S*_*β*_ than our simulations, possibly causing our estimates of the parameters to be upwardly biased. For this reason (and because the estimates of parameters are noisy in our simulations), we focus primarily on the direction of the effects rather than their magnitudes.

## Empirical data analyses

### Aggregating phenotype data

We obtained GWAS summary data from several published studies for nine different phenotypes as discussed in the main text (Wood et al., 2014; Shungin et al., 2015; Locke et al., 2015; CDG Psychiatric Genomics Consortium, 2013; Franke et al., 2010; Day et al., 2015; Global Lipids Genetics Consortium et al., 2013; Okbay et al., 2016). We obtained allele frequencies for each allele in each study from the GWAS summary data in the corresponding study. To calculate *S*_*β*_ for each study, we first needed to polarize all the alleles by their derived/ancestral status. We obtained inferred derived/ancestral states for each allele from the 1000 Genomes project (Thousand Genomes Project Consortium et al., 2015). Our permutation method requires that each allele then be assigned to a LD block within the genome (Berisa and Pickrell, 2016). To assign each of these alleles to LD blocks, we also used the 1000 Genomes data to obtain the genomic coordinates for each allele. Alleles for which we could not assign states or LD blocks were excluded.

### Calculating *S*_*β*_ and implementing our empirical framework

To account for LD in selection tests for complex traits using GWAS summary data, we divided the genome into 1,703 LD blocks, which were previously identified as being approximately independent (Berisa and Pickrell, 2016). For each LD block, we then select a random sign (positive or negative with equal probability), and multiply all the effect sizes in the LD block by this sign. We then recompute summary statistics (such as the correlation between frequency and MAF) on the randomized data. By repeating this procedure, we generate a null distribution for the test statistic. This method maintains the correlations between effect sizes generated by LD, the site frequency spectrum of the sampled alleles, and the joint distribution of the absolute value of effect size and allele frequency, while breaking any relationship between effect sizes and allele frequency. Note that this is a conservative permutation, because many of the alleles within an LD block are not linked or only weakly linked. We further consider the robustness of our method to population stratification in a subsequent section, which is a persistent potential source of false positives for studies of selection. To assess significance, we perform a two-tailed test comparing the observed value of the test statistic to the permutation-based null distribution. We performed 2,000 permutations for each phenotype.

We developed custom software implementing our approach in Python, which is freely available upon request and will be posted on Github. To calculate *S*_*β*_, we group alleles into 1% frequency bins – without performing this grouping, many high frequency bins would have very few alleles and very noisy estimates of 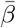. We selected 1% because it strikes a balance between obtaining low standard errors on 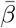 and finely parsing the allele frequency space, but we note that other choices of bin size could potentially improve our power.

### Replication data & analysis

As a replication cohort for our top three selection signals, we obtained GWAS summary statistics from the UK Biobank (UKBB) from https://data.broadinstitute.org/alkesgroup/UKBB/ (height, BMI, and educational attainment – note that approximately ¼ of the UKBB samples were included in Okbay *et al*, so the studies are not completely independent). We used our framework to calculate *S*_*β*_ on each phenotype, and estimate the linkage-adjusted variance in the test-statistic. We perform a one-tailed test for significance, since we have an expectation for the sign of *S*_*β*_ from our first set of findings. We replicate our findings for both BMI (*p* = 0.0095) and educational attainment (*p* < 5*e* − 4) (Fig. S18). Perhaps surprisingly, height did not replicate (see main text for discussion of possible explanations).

In addition to *S*_*β*_, we performed a replication study at signals of selection that were used in previous projects to identify signals of selection on height in the GIANT study. One study identified a positive correlation between effect size and allele frequency (Yang et al., 2015), while another found a correlation of allele frequency difference between northern and southern Europe and the p-value rank of the height increasing-allele (Turchin et al., 2012). Neither of these correlations were replicated in the UK Biobank summary statistics (Fig. S6).

### Neanderthal alleles

A list of Neanderthal alleles that were inferred with the *S*^*∗*^ statistic (Plagnol and Wall, 2006) was generously provided by Rajiv McCoy and Josh Akey. In analyses using Neanderthal alleles, we masked all alleles that were not present in this dataset. The dataset includes 139,694 SNPs, 139,381 of which we were able to map to rs numbers and use in our analysis.

*Neanderthal polarization and alternate summary statistics*. Although we have highlighted the ancestral/derived polarization, other polarizations, such as Neanderthal vs ancestral human alleles, can also provide insight into evolutionary processes. Recent evidence suggests that Neanderthal immune-related variants and expression-altering variants were likely targets of selection (Quach et al., 2016; Nédélec et al., 2016; McCoy et al., 2017), and some Neanderthal variants are likely to alter complex traits (Simonti et al., 2016), but little is known about polygenic selection on trait-altering Neanderthal alleles. If Neanderthals were “pre-adapted” to Europe, alleles that were fixed in Neanderthals may have accelerated the adaptation of modern humans to the European environment when admixed into humans (Enard and Petrov, 2017). Hence, we hypothesized that we may observe signals in *S*_*β*_ if alleles that fixed on the Neanderthal lineage were systematically biased in one direction than the other, or if selection acted to promote or remove Neanderthal alleles with specific phenotypic effects once admixed into modern humans.

We tested this hypothesis by computing *S*_*β*_ on Neanderthal alleles that were included in previous GWAS. The alleles were identified with the *S*^*∗*^ statistic (Plagnol and Wall, 2006) and allele sharing with Neanderthal genomes, and hence do not suffer from the same potential biases from ancestral uncertainty as the derived/ancestral polarization. We performed both a common allele test (*S*_*β*_ (0.05,0.25)) and an all allele test (*S*_*β*_ (0,0.25)) for each of the nine phenotypes. We exclude alleles above frequency 0.25 because the vast majority of Neanderthal alleles have frequencies below 0.25.

Across the nine phenotypes we studied, we found signals for both height (*p* <5e-4) and major depression (*p* <5e-4) (Fig. S18), and a marginally significant signal for schizophrenia that did not pass a multiple testing correction (*p* =0.015). The height and schizophrenia signals were strongest when including only common alleles, whereas the depression signal was driven primarily by rare alleles. The height signal suggests selection that favored Neanderthal height-increasing alleles (either on the Neanderthal lineage or in modern humans), while the depression signal suggests that Neanderthal alleles had a large impact on depression risk, which have been preferentially pushed to low frequency by selection. The schizophrenia signal suggests selection against schizophrenia risk or a bias towards protective alleles from Neanderthals. No other phenotypes had significant p-values.

### The impact of ancestral uncertainty on *S*_*β*_

Ancestral states are not directly observed in genomic data, and are typically inferred by comparing human sequences to those of an out-group. The underlying assumption is that the allelic state in the out-group represents the most likely ancestral state for the allele. While this approach correctly assigns the ancestral state for the majority of alleles, it does not account for recurrent fixation events at a single site, leading to some rate of ancestral misassignment. Since our method to detect selection compares ancestral and derived alleles, we sought to understand how ancestral misassignment would impact our inferences.

Ancestral misassignment will tend to decrease the absolute value of *S*_*β*_ within each frequency bin, and hence decrease our power. To see this, suppose that for a given minor allele frequency *x*, we misassign ancestral state with probability 1 − *p*, and Ψ(*x*) is the number of derived alleles observed at frequency x. We note that we can rewrite *S*_*β*_ as the difference in mean beta between ancestral and derived alleles of equal minor allele frequency (*i.e.*, 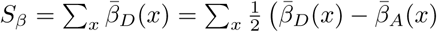). The expected value of the the test statistic within this bin is then

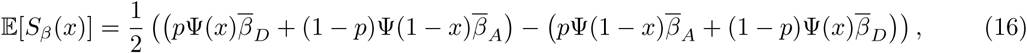

where *β*_*D*_ is the effect size of derived alleles and *β*_*A*_ is the effect size of ancestral alleles. Note that if *p* = 0.5, we are randomly guessing at ancestral states, and 𝔼[*S*_*β*_ (*x*)] = 0. If *p* < 0.5, then the absolute value of 𝔼[*S*_*β*_(*x*)] is strictly less than it’s true value (*i.e.*, the value that would be observed with no ancestral uncertainty).

While this analysis shows that ancestral uncertainty serves to make our analyses more conservative, we also sought to understand its potential impact on the observed relationship between allele frequency and effect size (Fig. S8). Because Ψ(*x*) is generally much larger than Ψ(1 − *x*) for *x* < 0.5 (*i.e.*, there are more rare alleles than common alleles in genomic sequencing data), we expect that the impact of ancestral uncertainty will be greatest at very high derived allele frequencies (*e.g.*, *x >* 0.9). We fit a linear model by regressing the mean effect size on allele frequency for the observed height data for derived alleles in the frequency range 40-60%, and extrapolated this curve out to 100% frequency, supposing that frequencies near 50% were only modestly impacted by ancestral uncertainty. We then simulated the impact of ancestral uncertainty at various levels from *p* = 1% to *p* = 10%. At *p* = 10% uncertainty we see a striking resemblance between our simulated data and the observed data. We conclude that moderate levels of ancestral uncertainty are likely responsible for the “*S*” shaped curve that we observe for many of our phenotypes.

**Figure S1:**
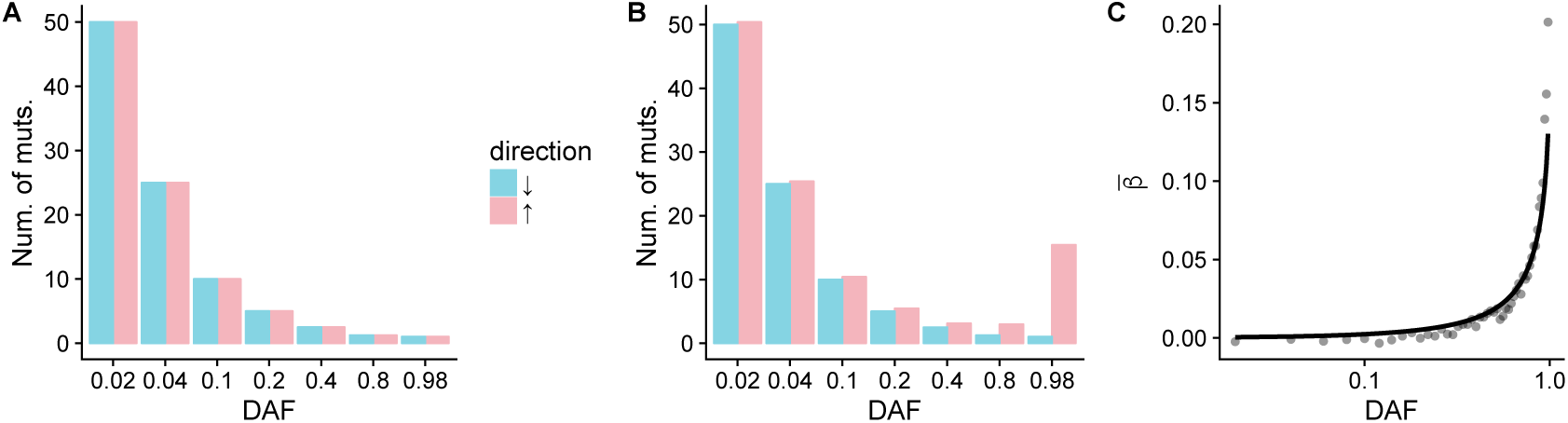
A: When selection does not prefer either trait-increasing or trait-decreasing alleles, and mutation bias does not act on a trait, then trait-increasing (pink) and - decreasing (blue) alleles are expected to have identical frequency spectra and equal mean *β* values within each frequency bin. B: When selection acts to increase a trait, trait-increasing alleles will increase in frequency and persist longer than trait-decreasing alleles, resulting in a relationship between 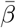 and frequency. C: 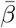 vs DAF for a toy model of where trait-increasing alleles are favored. The black line represents an analytical calculation (eqn. 5) while the black points are the results of stochastic simulations.

**Figure S2:**
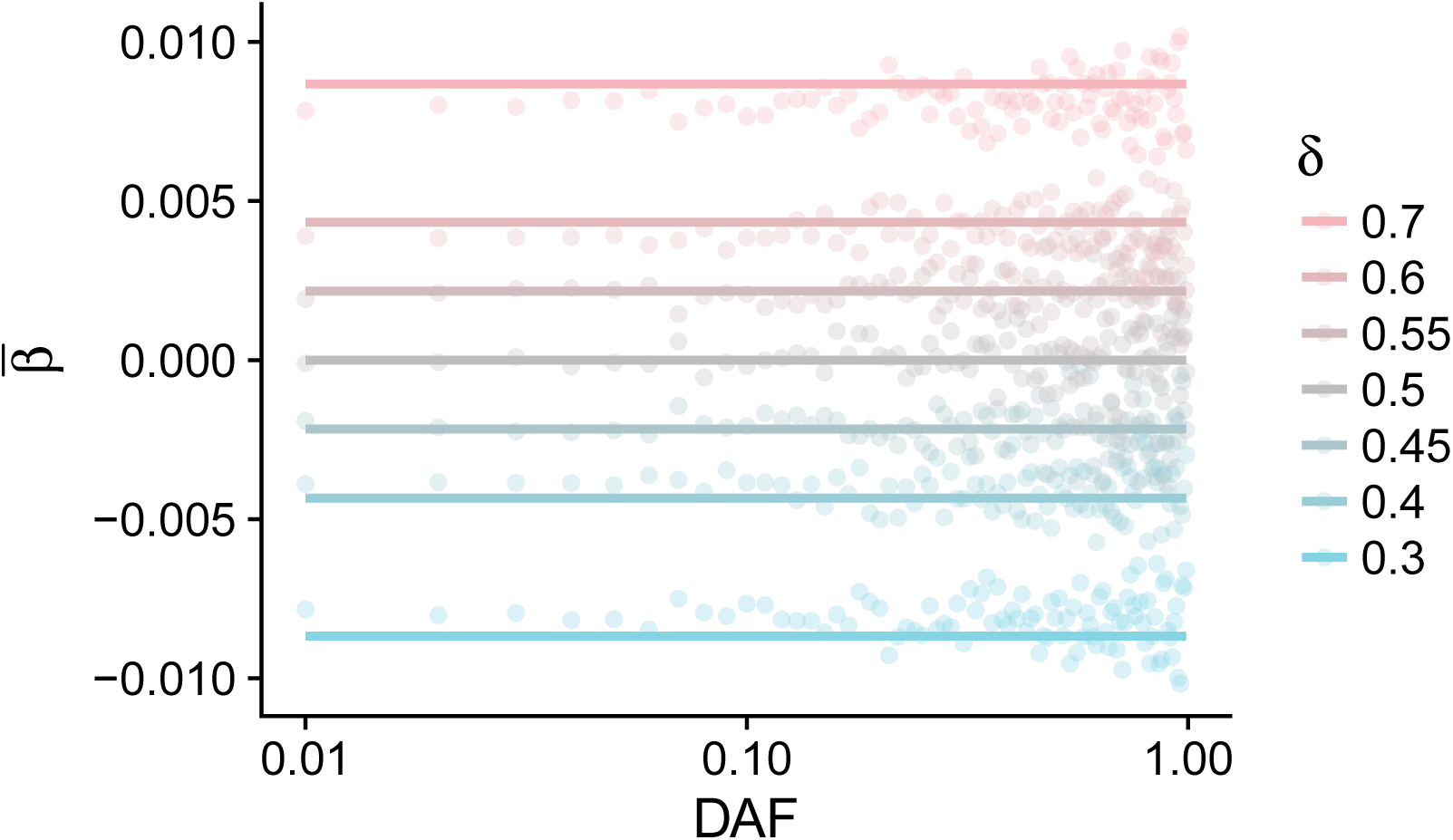
The relationship between *β* and DAF when only mutational bias acts on the trait (*i.e.*, there is no selection). Mean *β* is shifted in the direction of the bias, but does not depend on DAF.

**Figure S3:**
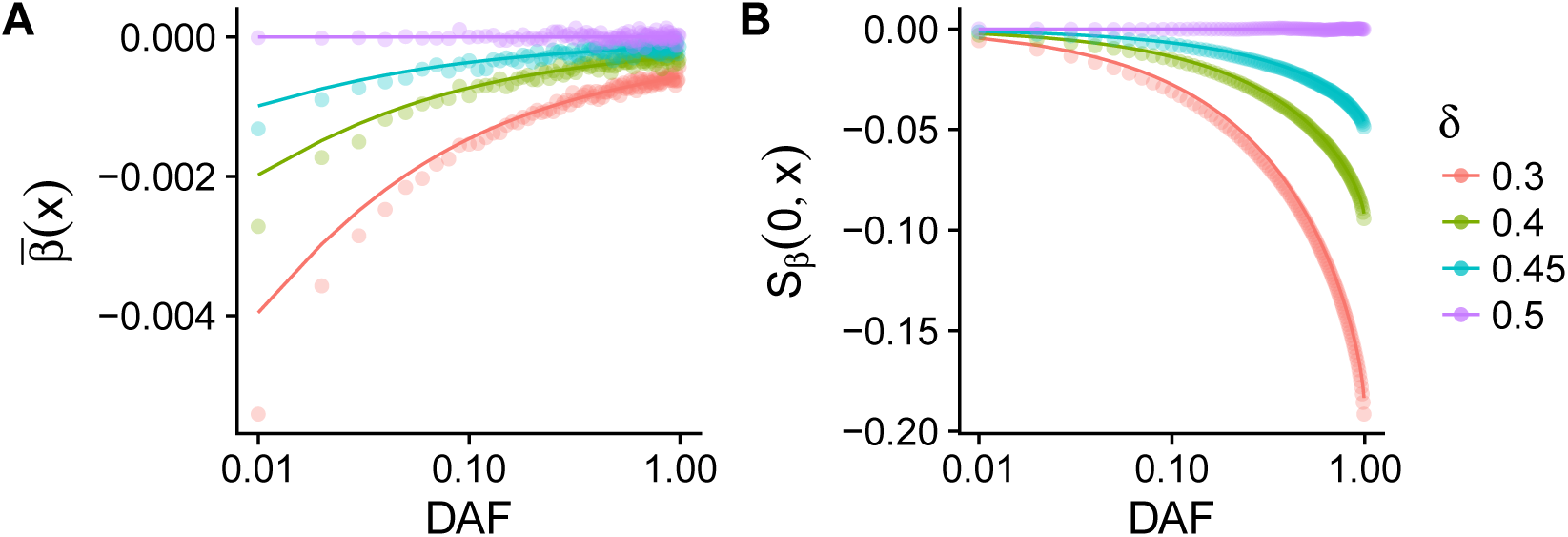
A: A *β*DAF plot for various values of mutational bias (*δ*) towards trait decreasing alleles for traits under selection. B: *S*_*β*_(0, *x*) for the same models in A. The solid lines represent the results of analytical calculations (eqn. 6) while the points represent stochastic simulations.

**Figure S4:**
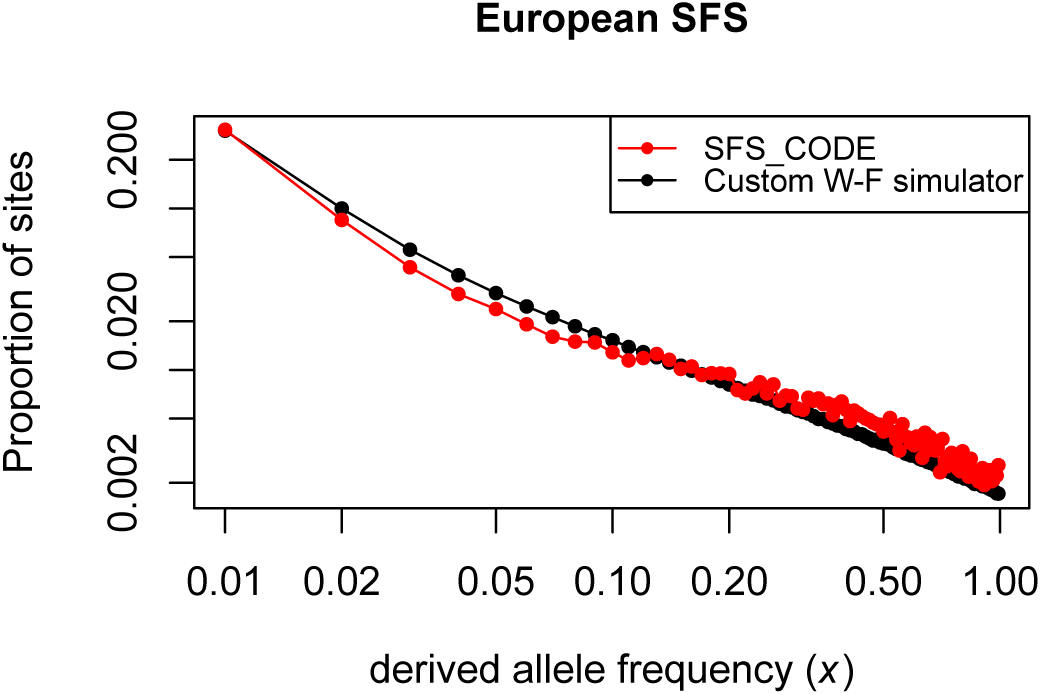
A comparison of simulated frequency spectra for a complex demographic/selection model for our Wright-Fisher simulator (black) and SFS CODE (red).

**Figure S5:**
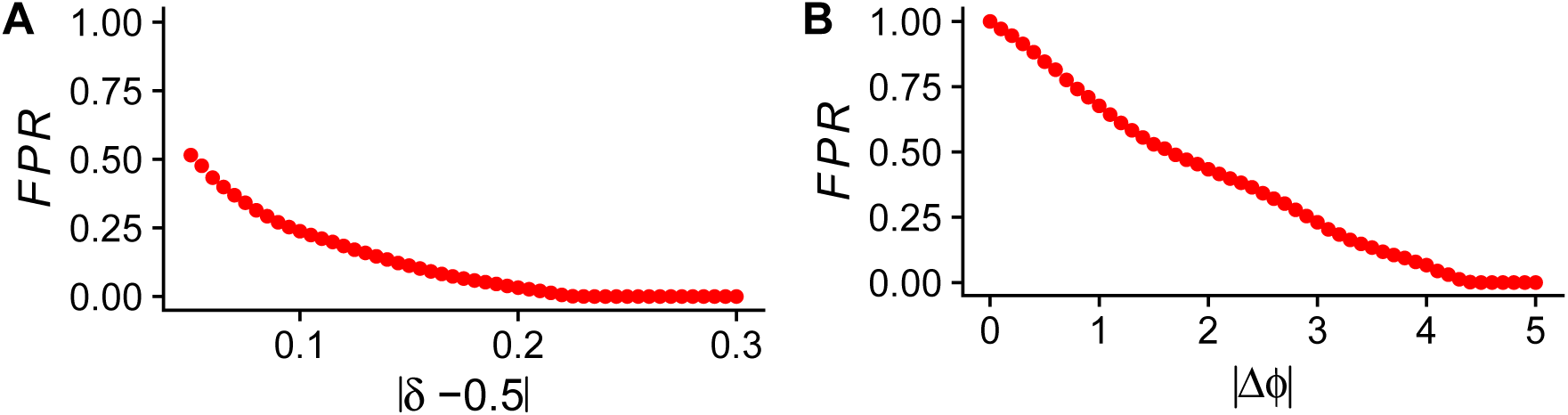
Conservative False Positive Rate estimation for detection of mutation rate bias and shifts in optimal phenotype value. We plot the estimated probability (FPR) that the inferred parameter exceeds the value on the x-axis, given that no mutation bias (A) or shift in optimal phenotype (B) occurred. As expected, it is not unlikely to infer a small shift in cases when no shift occurred, but it is highly unlikely to infer a large shift.

**Figure S6:**
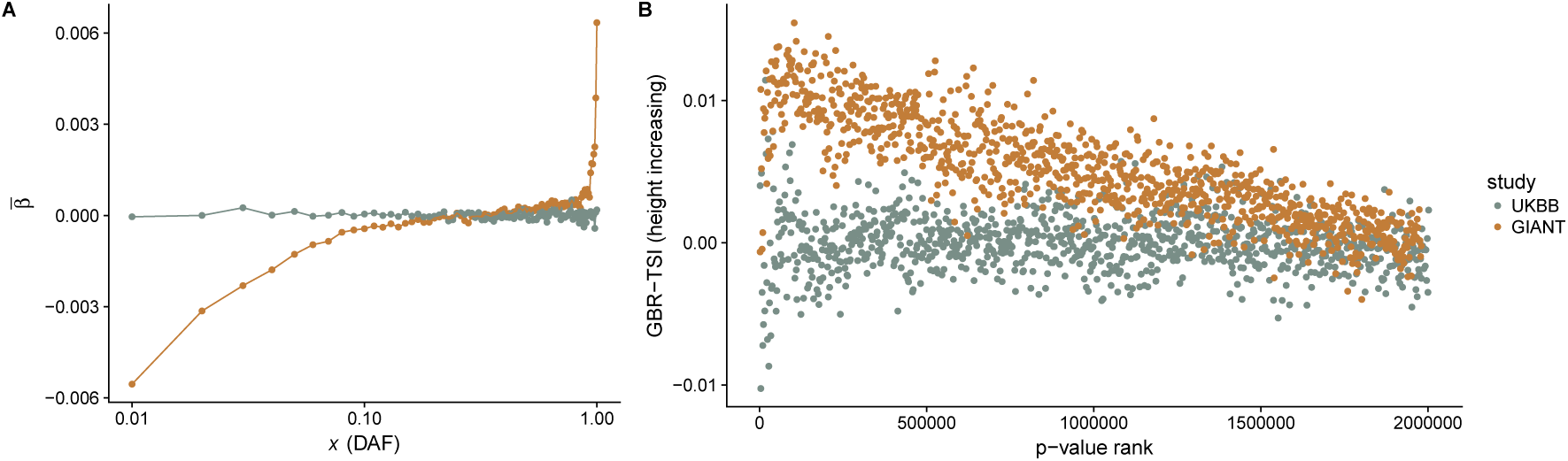
Correlation between mean effect size and allele frequency (A) and correlation the difference in frequency between northern and southern Europe for the height increasing allele and p-value rank (B) in the GIANT study (gold) and the UK Biobank (gray). Note that alleles were grouped in to bins of 2,000 by p-value and were not thinned by linkage.

**Figure S7:**
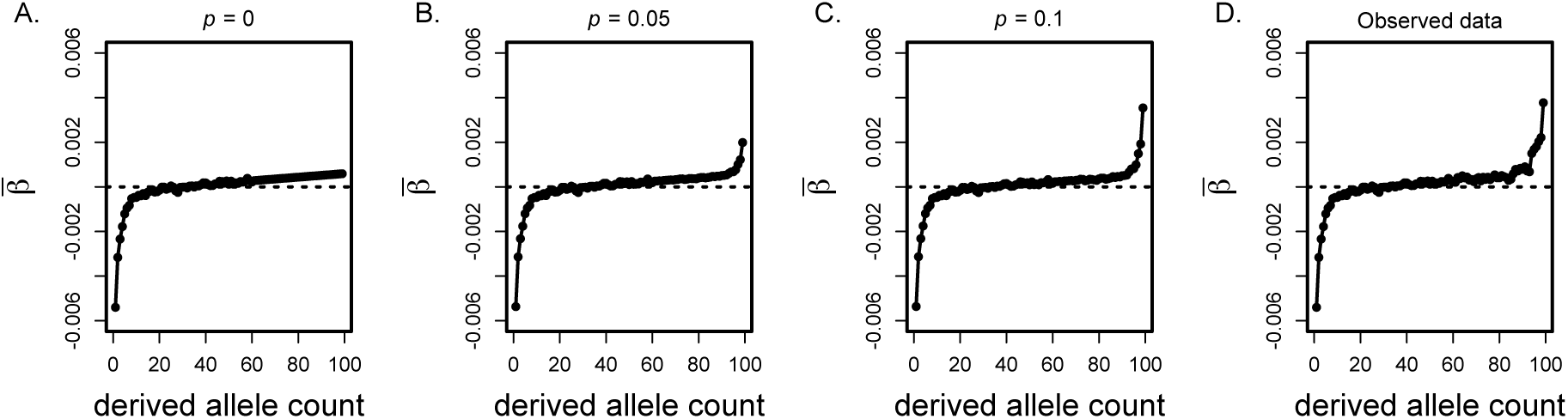
The impact of ancestral uncertainty on the observed value of mean *β* as a function of derived allele count *x*. Panel A shows results corresponding to no uncertainty (*p* = 0), in B *p* = 0.05, in C *p* = 0.1, and D shows the observed GIANT height data.

**Figure S8:**
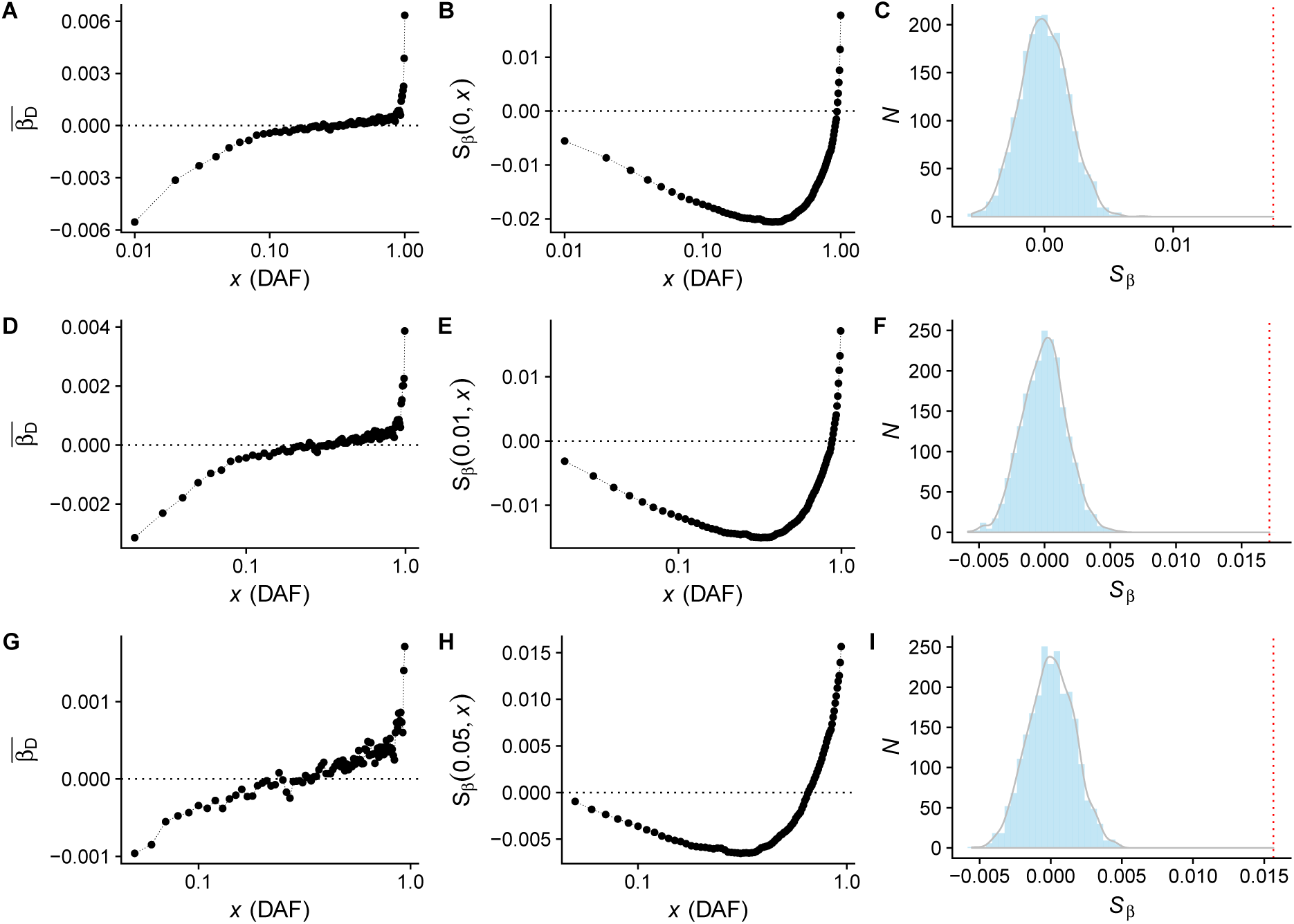
*S*_*β*_ for height. The panels in the left column show the relationship between allele frequency and 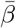, the middle column displays the cumulative value of *S*_*β*_(*x*_*i*_, *x*_*f*_), and the right columns show the null distribution of *S*_*β*_ given by our permutation test. Panels A-C correspond to *x*_*i*_ = 0 and *x*_*f*_ = 1, panels D-F correspond to *x*_*i*_ = 0.01 and *x*_*f*_ = 0.99, and panels G-I correspond to *x*_*i*_ = 0.05 and *x*_*f*_ = 0.95

**Figure S9:**
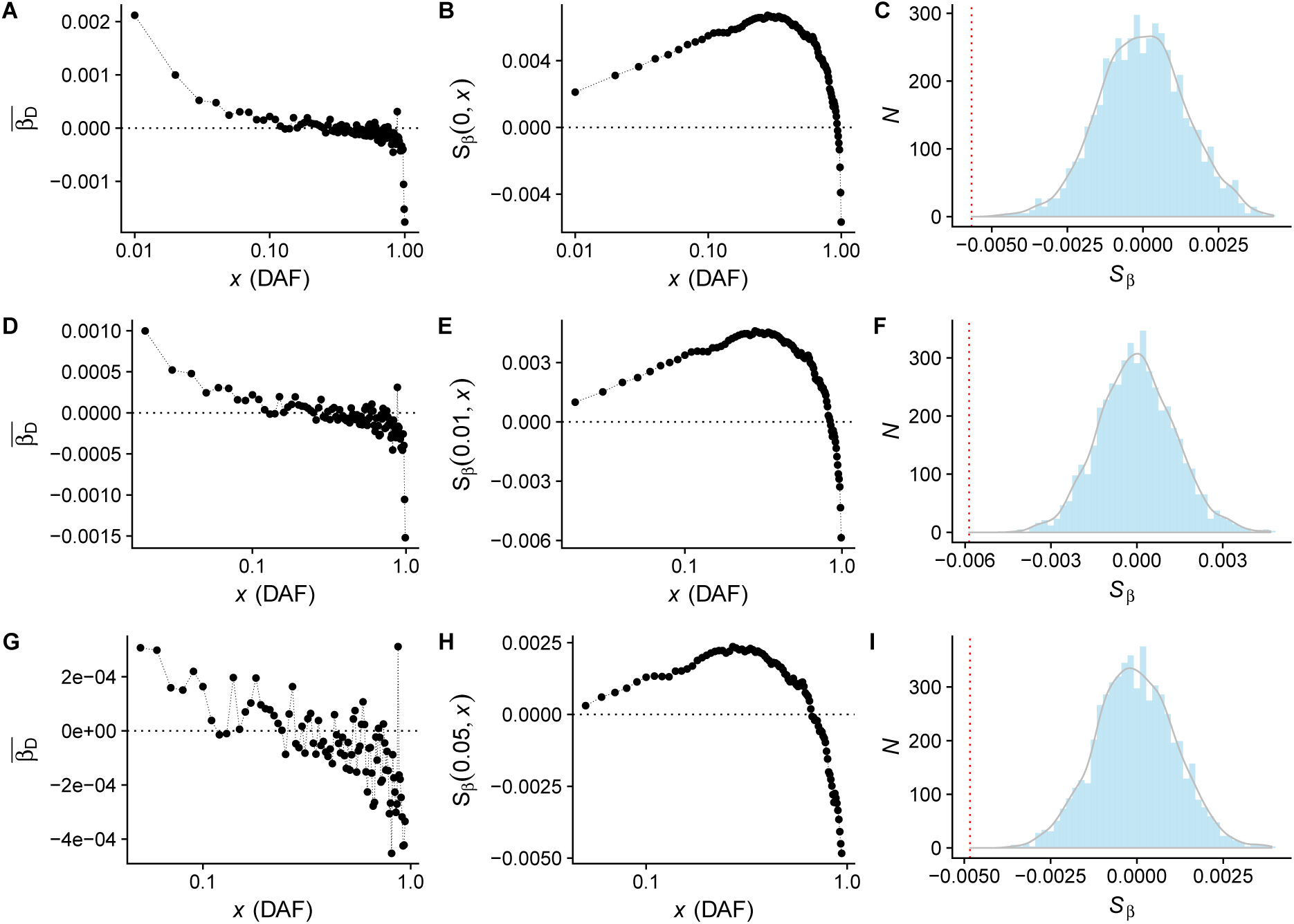
*S*_*β*_ for BMI. The panels in the left column show the relationship between allele frequency and 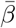, the middle column displays the cumulative value of *S*_*β*_(*x*_*i*_, *x*_*f*_), and the right columns show the null distribution of *S*_*β*_ given by our permutation test. Panels A-C correspond to *x*_*i*_ = 0 and *x*_*f*_ = 1, panels D-F correspond to *x*_*i*_ = 0.01 and *x*_*f*_ = 0.99, and panels G-I correspond to *x*_*i*_ = 0.05 and *x*_*f*_ = 0.95

**Figure S10:**
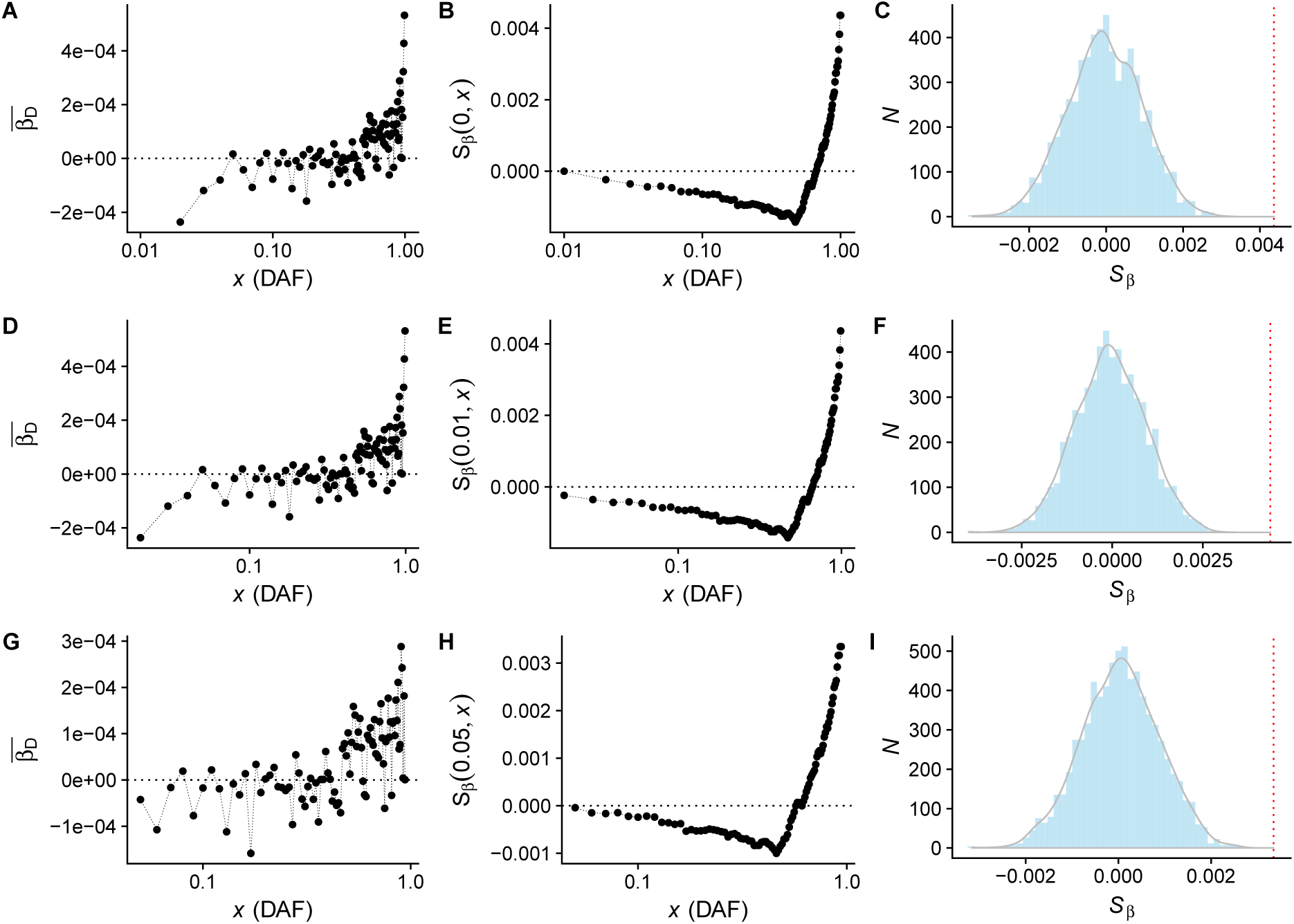
*S*_*β*_ for educational attainment. The panels in the left column show the relationship between allele frequency and 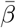, the middle column displays the cumulative value of *S*_*β*_(*x*_*i*_, *x*_*f*_), and the right columns show the null distribution of *S*_*β*_ given by our permutation test. Panels A-C correspond to *x*_*i*_ = 0 and *x*_*f*_ = 1, panels D-F correspond to *x*_*i*_ = 0.01 and *x*_*f*_ = 0.99, and panels G-I correspond to *x*_*i*_ = 0.05 and *x*_*f*_ = 0.95. Note that the results for A-C are the same as D-F because no alleles under 1% were included in the summary data for this study.

**Figure S11:**
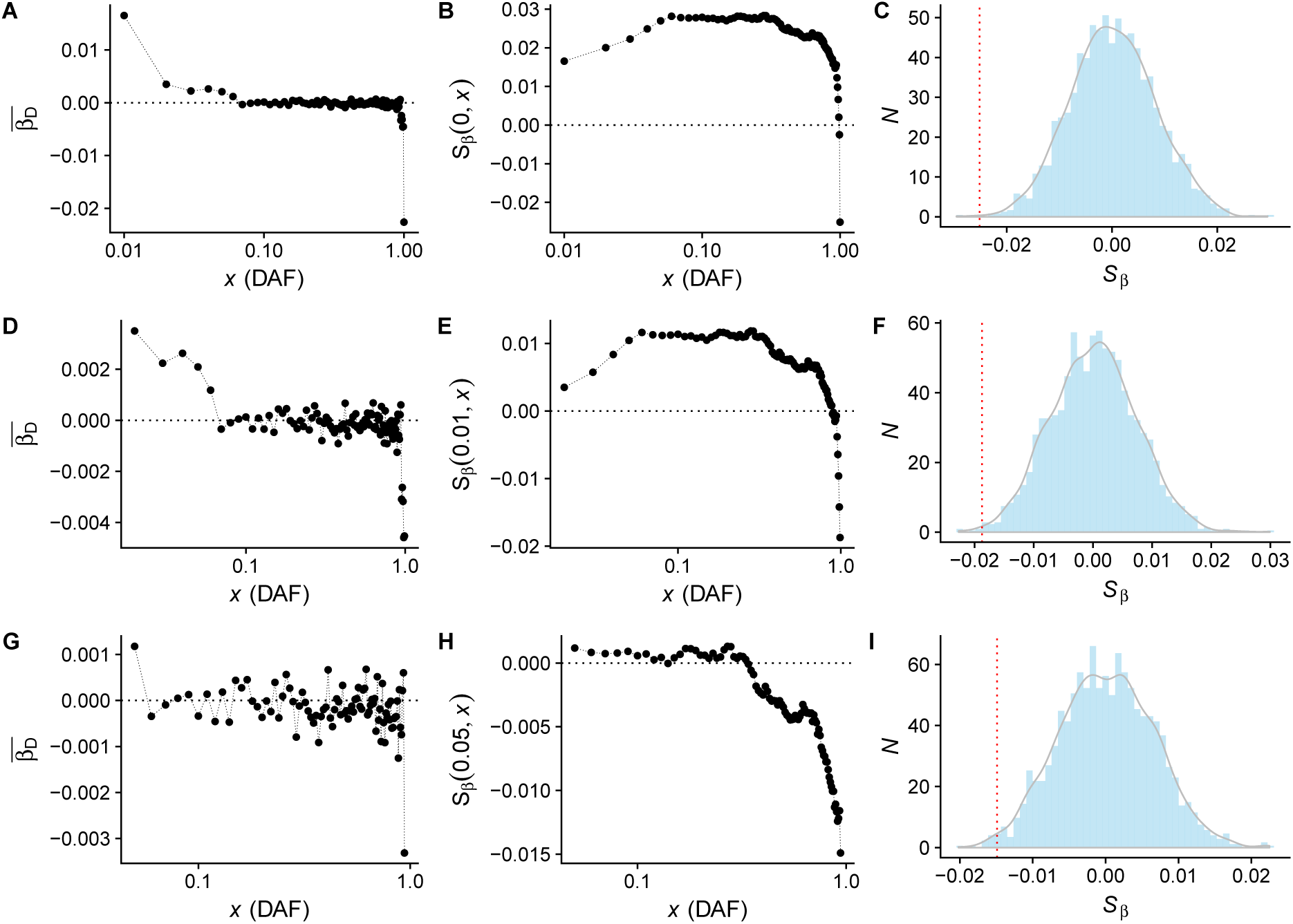
*S*_*β*_ for Crohn’s disease. The panels in the left column show the relationship between allele frequency and 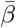, the middle column displays the cumulative value of *S*_*β*_(*x*_*i*_, *x*_*f*_), and the right columns show the null distribution of *S*_*β*_ given by our permutation test. Panels A-C correspond to *x*_*i*_ = 0 and *x*_*f*_ = 1, panels D-F correspond to *x*_*i*_ = 0.01 and *x*_*f*_ = 0.99, and panels G-I correspond to *x*_*i*_ = 0.05 and *x*_*f*_ = 0.95

**Figure S12:**
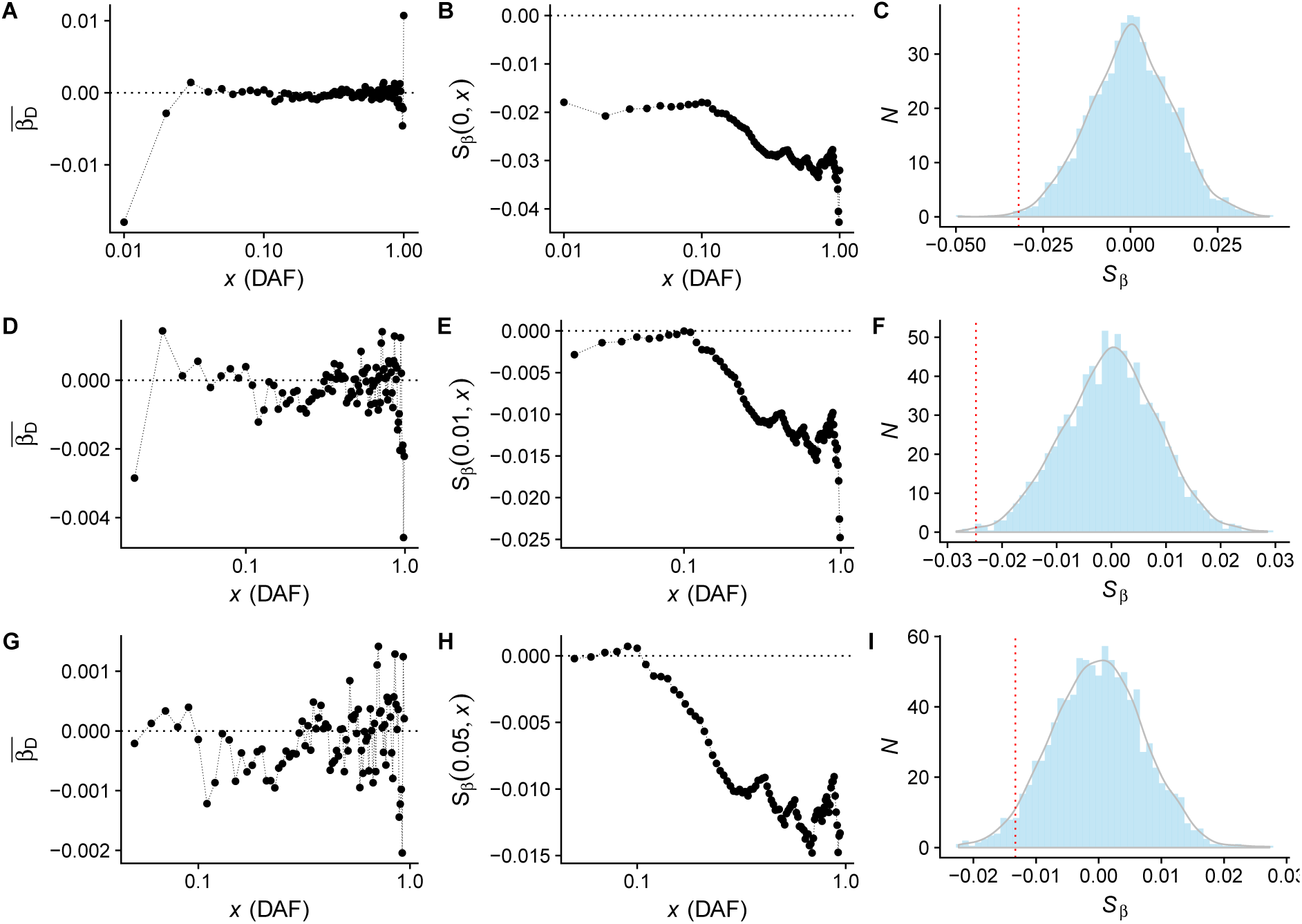
*S*_*β*_ for schizophrenia. The panels in the left column show the relationship between allele frequency and 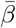, the middle column displays the cumulative value of *S*_*β*_(*x*_*i*_, *x*_*f*_), and the right columns show the null distribution of *S*_*β*_ given by our permutation test. Panels A-C correspond to *x*_*i*_ = 0 and *x*_*f*_ = 1, panels D-F correspond to *x*_*i*_ = 0.01 and *x*_*f*_ = 0.99, and panels G-I correspond to *x*_*i*_ = 0.05 and *x*_*f*_ = 0.95

**Figure S13:**
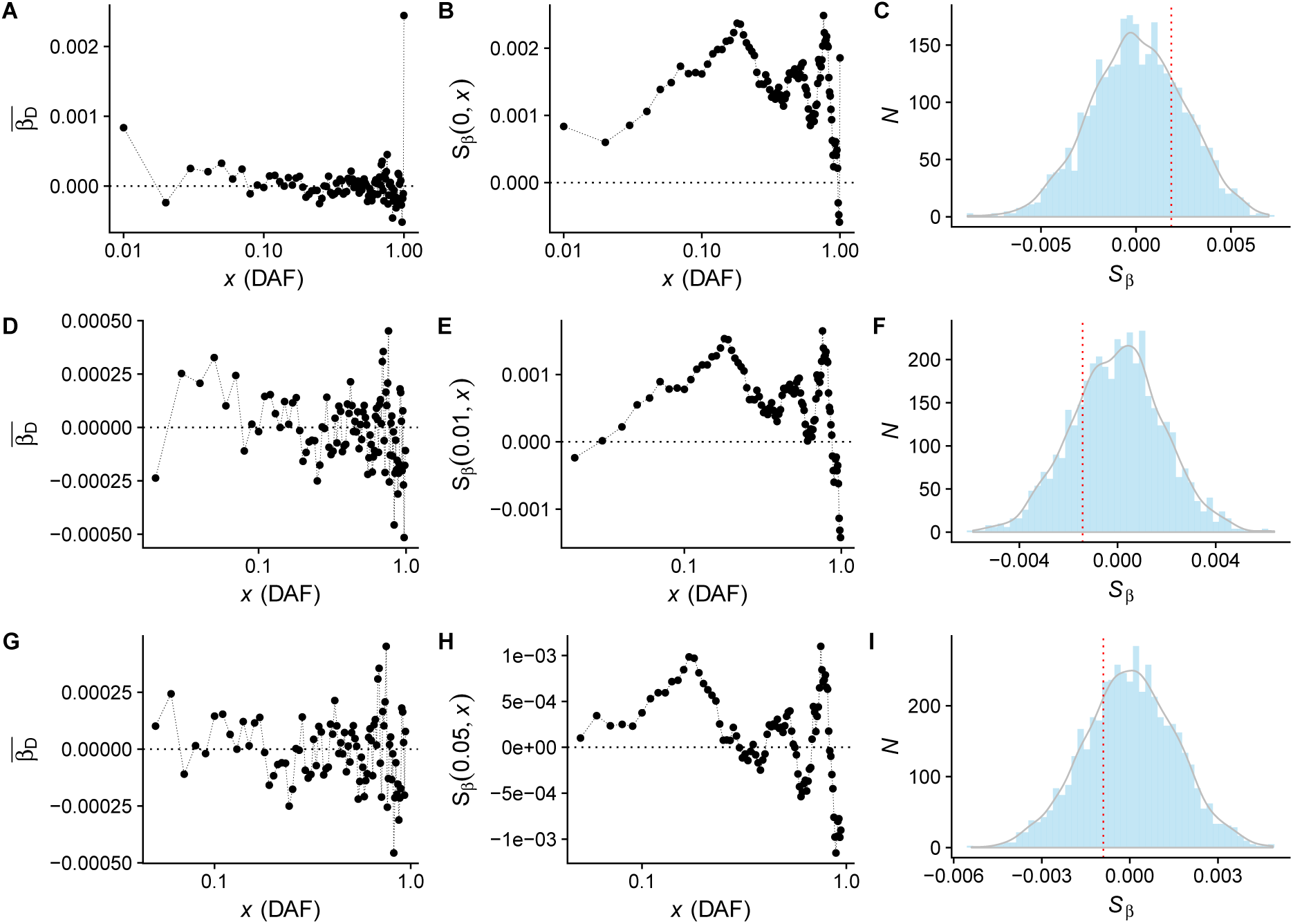
*S*_*β*_ for global lipid levels. The panels in the left column show the relationship between allele frequency and 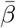, the middle column displays the cumulative value of *S*_*β*_(*x*_*i*_, *x*_*f*_), and the right columns show the null distribution of *S*_*β*_ given by our permutation test. Panels A-C correspond to *x*_*i*_ = 0 and *x*_*f*_ = 1, panels D-F correspond to *x*_*i*_ = 0.01 and *x*_*f*_ = 0.99, and panels G-I correspond to *x*_*i*_ = 0.05 and *x*_*f*_ = 0.95

**Figure S14:**
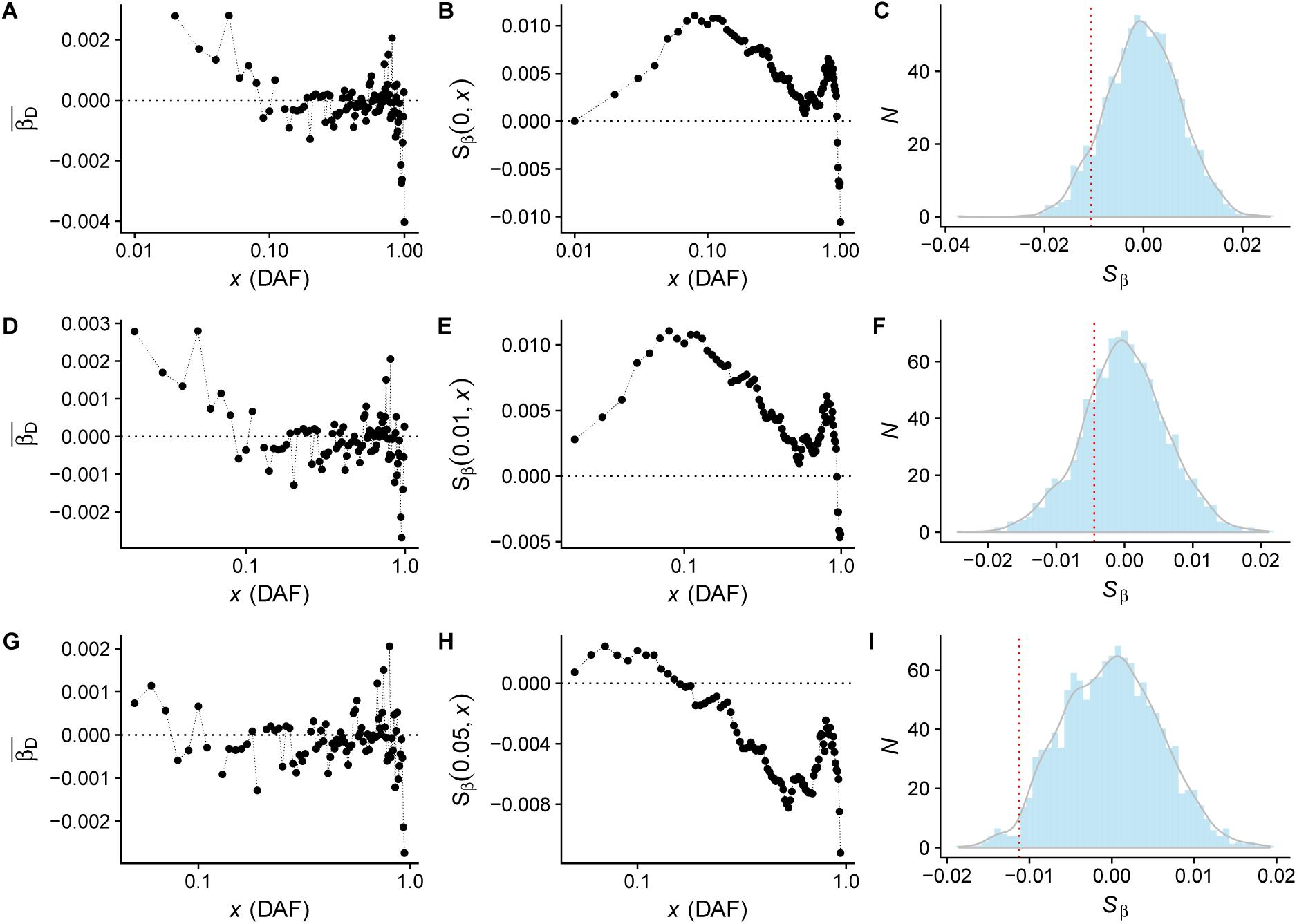
*S*_*β*_ for menopause onset. The panels in the left column show the relationship between allele frequency and 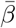, the middle column displays the cumulative value of *S*_*β*_(*x*_*i*_, *x*_*f*_), and the right columns show the null distribution of *S*_*β*_ given by our permutation test. Panels A-C correspond to *x*_*i*_ = 0 and *x*_*f*_ = 1, panels D-F correspond to *x*_*i*_ = 0.01 and *x*_*f*_ = 0.99, and panels G-I correspond to *x*_*i*_ = 0.05 and *x*_*f*_ = 0.95

**Figure S15:**
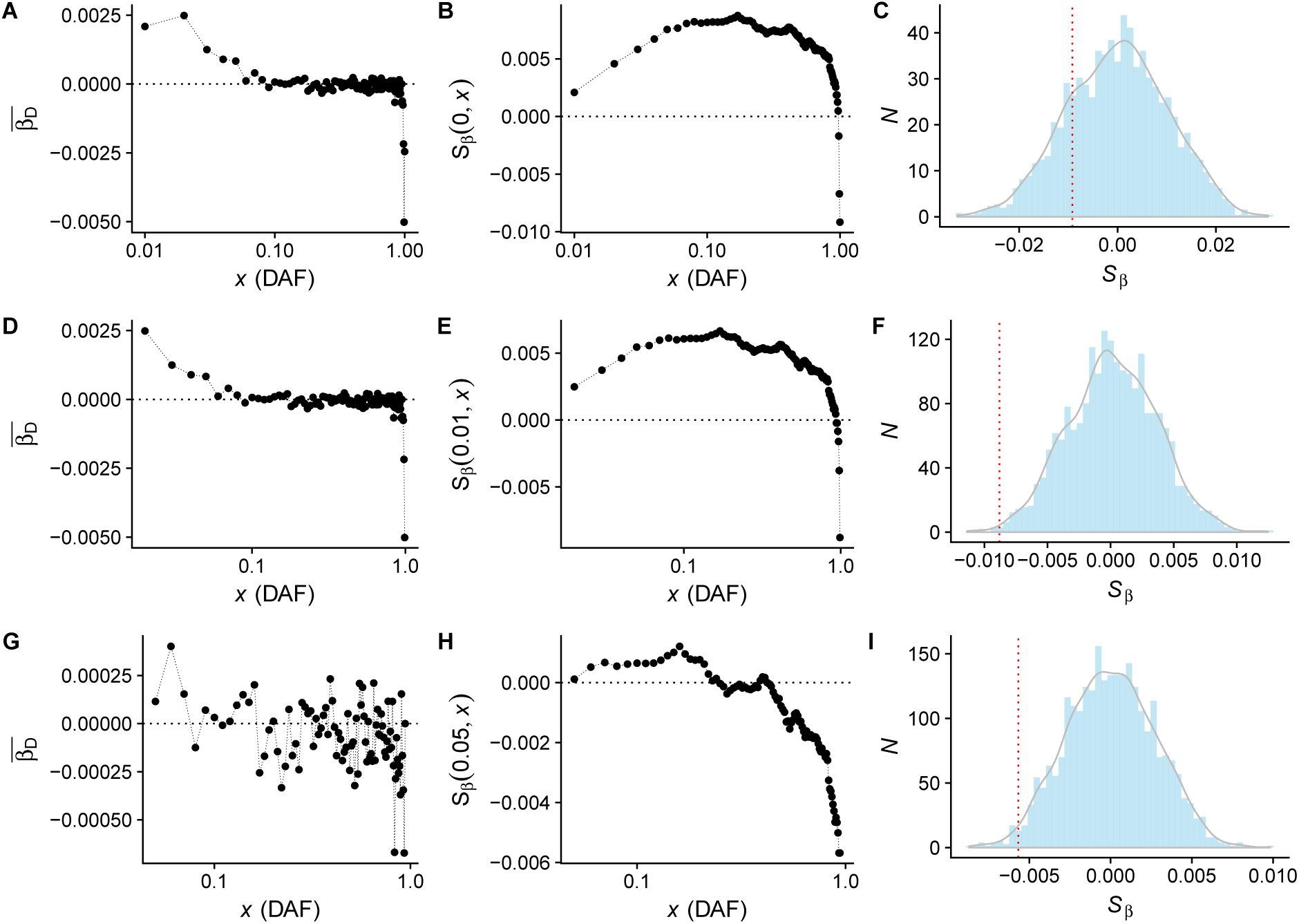
*S*_*β*_ for major depression. The panels in the left column show the relationship between allele frequency and 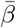, the middle column displays the cumulative value of *S*_*β*_(*x*_*i*_, *x*_*f*_), and the right columns show the null distribution of *S*_*β*_ given by our permutation test. Panels A-C correspond to *x*_*i*_ = 0 and *x*_*f*_ = 1, panels D-F correspond to *x*_*i*_ = 0.01 and *x*_*f*_ = 0.99, and panels G-I correspond to *x*_*i*_ = 0.05 and *x*_*f*_ = 0.95

**Figure S16:**
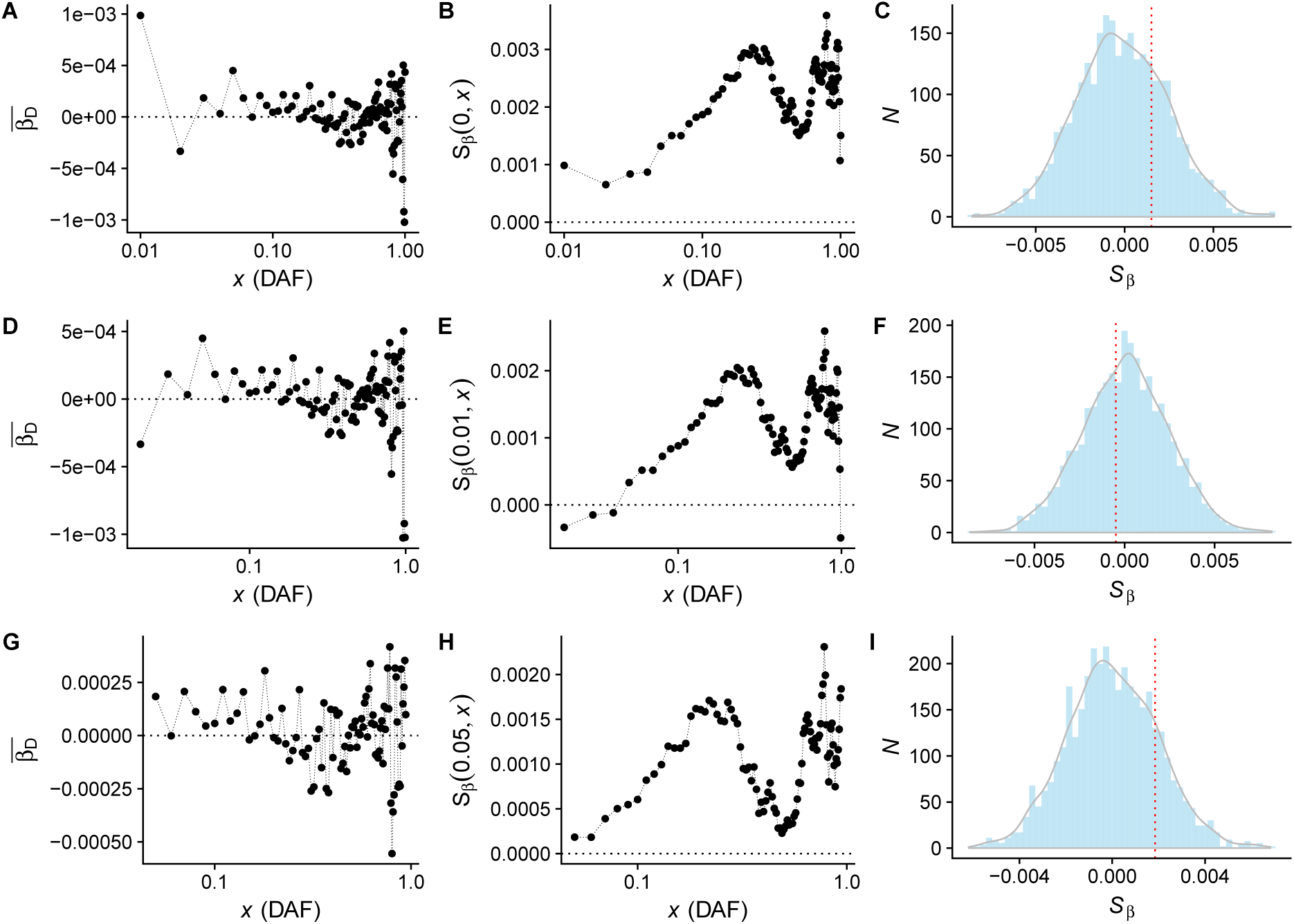
*S*_*β*_ for waist-hip ratio adjusted for BMI. The panels in the left column show the relationship between allele frequency and 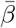, the middle column displays the cumulative value of *S*_*β*_(*x*_*i*_, *x*_*f*_), and the right columns show the null distribution of *S*_*β*_ given by our permutation framework. Panels A-C correspond to *x*_*i*_ = 0 and *x*_*f*_ = 1, panels D-F correspond to *x*_*i*_ = 0.01 and *x*_*f*_ = 0.99, and panels G-I correspond to *x*_*i*_ = 0.05 and *x*_*f*_ = 0.95

**Figure S17:**
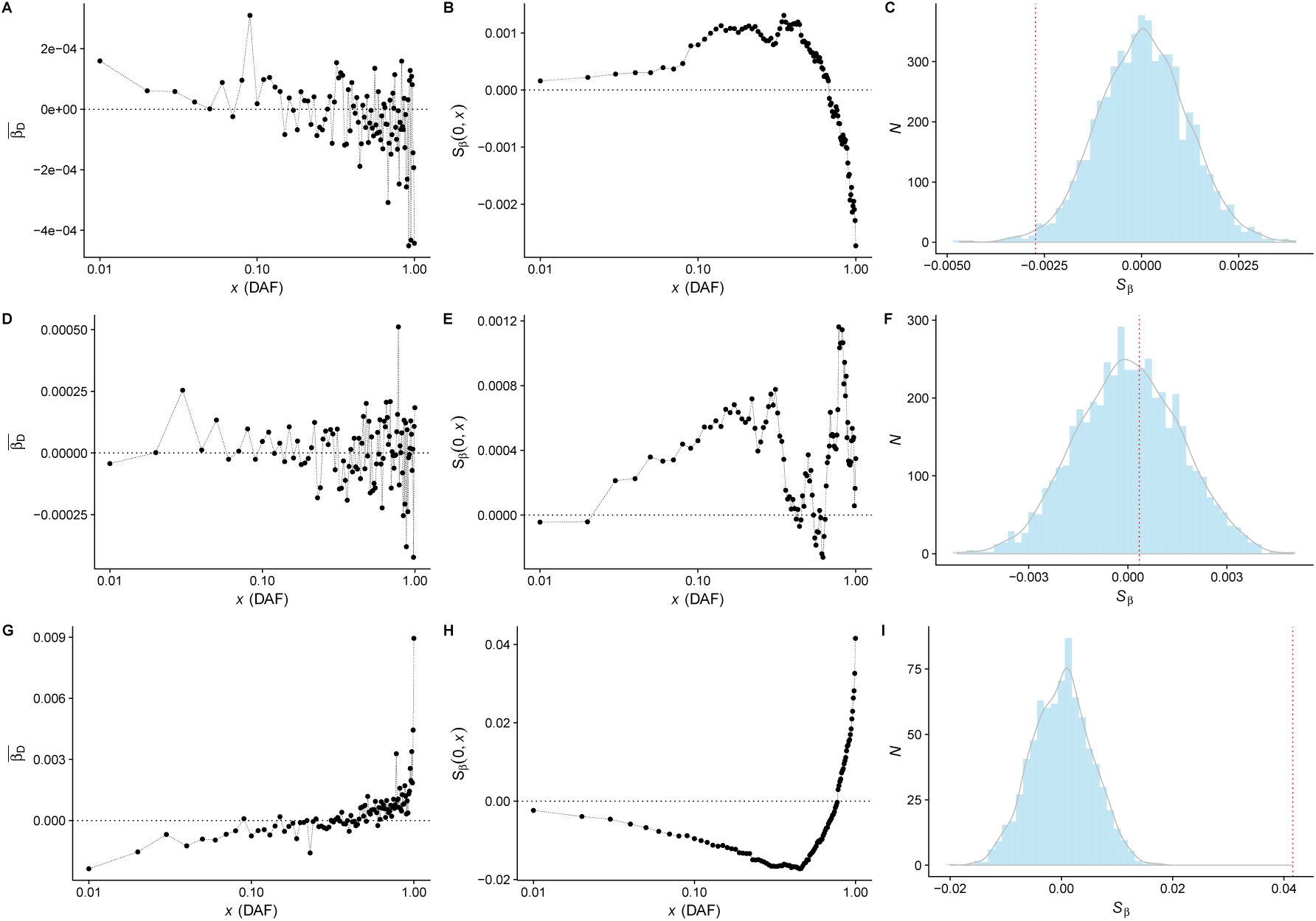
*S*_*β*_ for BMI (A-C), height (D-F), and educational attainment (G-I) in the UK Biobank. The panels in the left column show the relationship between allele frequency and 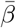, the middle column displays the cumulative value of *S*_*β*_(*x*_*i*_, *x*_*f*_), and the right columns show the null distribution of *S*_*β*_ given by our permutation framework.

**Figure S18:**
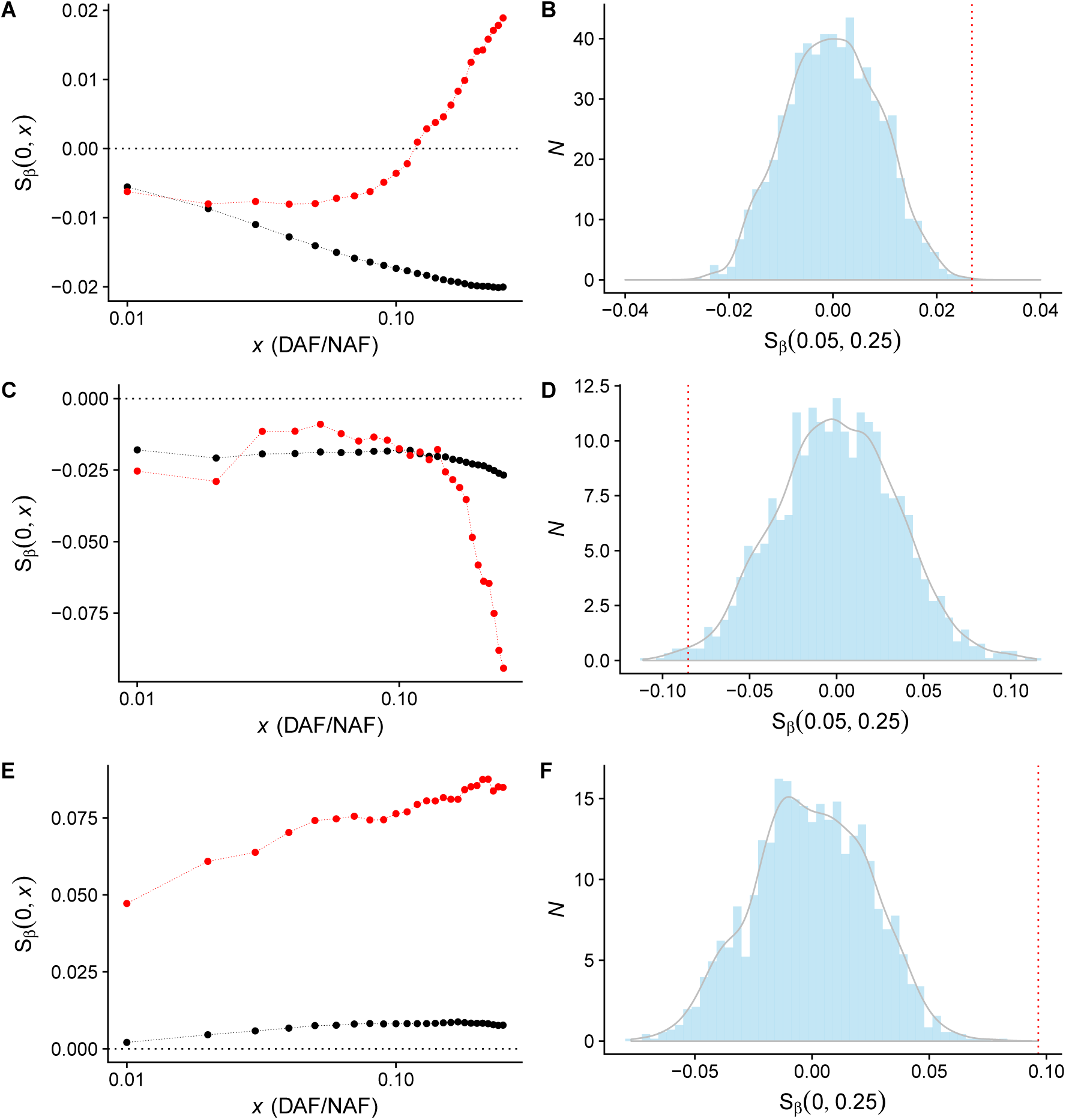
Signals of putative polygenic selection and mutation bias on Neanderthal alleles. A-B correspond to height, C-D correspond to schizophrenia, and E-F correspond to major depression. The left column shows *S*_*β*_(0, *x*) for Neanderthal alleles (red) and modern human alleles for comparison (truncated at allele frequency of 0.25). The right column shows *S*_*β*_(*x*_*i*_ 0.25) for our permutations, where the observed signal is shown in the red dashed line and the value of *x*_*i*_ is indicated under the plot. We note that

**Figure S19:**
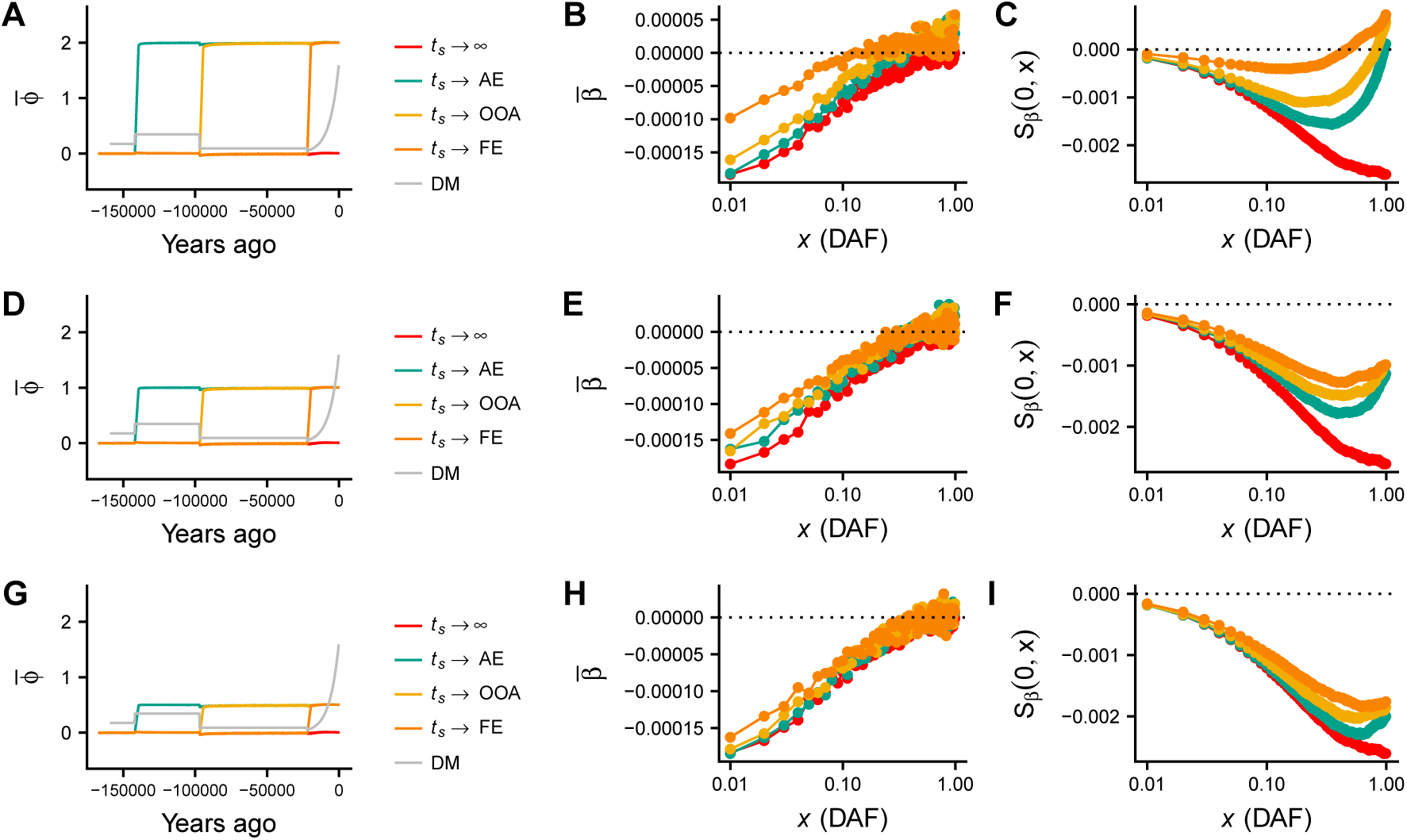
A-C: Simulations with the same parameters as Fig. 1C-E, but with the shift in optimal phenotype of ∆*ϕ* = 2 occurring linearly over 100 generations (2500 years), rather than instantaneously in a single generation. D-F: Simulations with the same parameters as Fig. 1C-E, but with a shift of ∆*ϕ* = 1 occurring linearly over 100 generations. G-I: Simulations with the same parameters as Fig. 1C-E, but with a shift of ∆*ϕ* = 0.5 occurring linearly over 100 generations.

